# Multiple parallel expansions of bilaterian-like phototransduction gene families in the eyeless Anthozoa

**DOI:** 10.1101/2023.09.14.557824

**Authors:** Stacey Hansen, Meghan Payne, Kyle J McCulloch

## Abstract

Opsin-mediated phototransduction cascades in photoreceptor cells are primarily responsible for light-mediated behaviors in animals. Although some visual cascades are well-studied, phototransduction mediated by non-visual opsins and in non-model animal lineages are poorly characterized. In the Cnidaria (jellyfish, corals, sea anemones etc.), the sister group to Bilateria (vertebrates, arthropods, mollusks etc.), limited evidence suggests some overlap with bilaterian phototransduction. This raises the question of whether phototransduction pathways arose a single time early in animal evolution or if light signaling cascades have evolved multiple times. These evolutionary patterns remain obscured because almost nothing is known about phototransduction in a major group within Cnidaria, the eyeless Anthozoa (corals, sea anemones, sea pens etc.). To better understand whether bilaterian-like phototransduction could be present in Anthozoa, we phylogenetically characterized 63 genes in 12 protein families known to be crucial in two types of bilaterian phototransduction in the sea anemone *Nematostella vectensis*. Using high quality genomic data from *N. vectensis,* we took a candidate gene approach to find phototransduction genes and characterize their expression in development and regeneration. We found that *N. vectensis* possesses the core suite of proteins for both r-opsin and c-opsin mediated phototransduction. In addition, several new gene subfamilies were identified, particularly in the G protein subunits and TRP channels, and many were anthozoan-specific. We identified a novel G protein α subunit family, which we call GαVI, and characterized its expression in *N. vectensis* with *in situ* hybridization. This expansion of phototransduction genes correlates with a large anthozoan-specific radiation in opsin number, suggesting possible coevolution of receptor and signaling diversity in Anthozoa. While further functional experiments on these genes are needed, our findings are in line with the hypothesis that the common ancestor of Eumetazoa had at least two related phototransduction cascades which then further diversified in each animal lineage.

## 1. Introduction

Nearly all animals utilize opsin-based light-sensing in a wide variety of contexts for survival and reproduction (Terakita, 2005). This type of light sensing is mediated at the cellular level by specific G protein signaling cascades in photoreceptor cells, known as phototransduction cascades (Shichida and Matsuyama, 2009; Yau and Hardie, 2009). Opsins are part of the G protein-coupled receptor (GPCR) superfamily, and together with a vitamin A derived chromophore are termed rhodopsins (Okada et al., 2004; Palczewski et al., 2000). Upon absorbing a photon of light, the rhodopsin complex activates G protein signaling in the cell via its specific binding to a particular Gα subunit in the G protein complex (Simon et al., 1991; Yau and Hardie, 2009). There are at least 4 known pathways that can be defined by the opsin and G protein complex that initiate the cascade in a cell (Fain et al., 2010). The best studied cascades are associated with the ciliary opsins (c-opsins) and rhabdomeric opsins (r-opsins). C-opsins are best known in the context of vertebrate rods and cones, signaling through a specific G protein complex termed transducin (Gt), a phosphodiesterase (PDE), the second messenger cGMP, and cyclic nucleotide gated (CNG) channels (Lamb, 2013; Yau, 1994). In contrast, r-opsins are best known from microvillar regions of rhabdomeric photoreceptor cells in the compound eyes of insects (Wang and Montell, 2007). The r-opsin phototransduction cascade utilizes a Gαq class subunit, and signals through a phosphoinositide pathway involving phospholipase C (PLC) and transient receptor potential (TRP) channels (Hardie and Raghu, 2001; Katz and Minke, 2009; Wang and Montell, 2007).

While these pathways are elucidated in a limited number of contexts, phototransduction by canonical opsins in non-model organisms and by non-typical opsin groups is poorly understood (Arendt et al., 2004; Gornik et al., 2021; Hattar et al., 2003; Kozmik et al., 2008; Liegertová et al., 2015; Mason et al., 2012; Rawlinson et al., 2019; Velarde et al., 2005; Vöcking et al., 2022, 2017). This limits our understanding of the evolution of this pathway and visual systems more broadly. Was a single phototransduction cascade assembled once and was modified throughout animal evolution, or did multiple cascades *de novo* evolve independently to function in different contexts? It is thought that opsins evolved early in animals and that multiple classes of opsins were already present before the split of Bilateria and Cnidaria (Arendt et al., 2004; Ramirez et al., 2016; Schnitzler et al., 2012). Opsin photoreceptive functions have been conserved across animals, suggesting signaling cascades may also share an early animal origin. This is supported by investigating the “non-visual” tetraopsins, xenopsins and cnidopsins, where the limited data available suggest all or part of the cascade is similar to either r- or c-opsin cascades (Döring et al., 2020; Kayal et al., 2018; Kojima et al., 1997; Koyanagi et al., 2008; Kozmik et al., 2008; Liegertová et al., 2015; Plachetzki et al., 2012, 2010; Vöcking et al., 2017). To better understand the evolution of phototransduction cascades, we need to understand more about the pathways in animals outside of model bilaterians.

Cnidaria are sister to Bilateria and offer an important phylogenetic comparison to understand the evolution of complexity in animals. The few comparative studies on phototransduction in Cnidaria have almost exclusively focused on jellyfish and *Hydra* (Medusozoa) (Ekström et al., 2008; Plachetzki et al., 2010; Vöcking et al., 2022) Evidence in box jellies suggests that a Gαs subunit signals via the non-canonical AC enzyme, similar to bilaterian tetraopsin signaling (Koyanagi et al., 2008). In addition, cnidarians with eyes share many of the same molecular components as bilaterian eyes, such as developmental genes, phototransduction cascade members, and opsins (Kozmik et al., 2008). These similarities suggest that the genetic components could have assembled in a signaling pathway and associated with opsins before the split of Bilateria and Cnidaria. Conversely the mechanisms of light sensing in the other major cnidarian group, the eyeless Anthozoa, have received relatively little attention (Mason et al., 2012). A more complete view of anthozoan phototransduction will provide important comparative data with both Medusozoa and Bilateria, and could better elucidate how and when different cascades arose.

The emerging model sea anemone *Nematostella vectensis* is an anthozoan cnidarian that lives in shallow estuaries of brackish water and salt marshes (Layden et al., 2016). Development proceeds from egg to free-swimming larval stage known as the planula by about 48 hours at room temperature, then undergoes metamorphosis into a juvenile primary polyp with four tentacles after roughly ten days (Layden et al., 2016). The primary polyp grows in a nutrient dependent matter and is sexually mature by about 3 months. *N. vectensis* can also reproduce asexually via transverse fission and has robust adult whole-body regeneration. Mature *N. vectensis* adults have two germ layers and a mouth that is surrounded by up to sixteen tentacles (Layden et al., 2016). Rather than specialized organs like an eye, specialized cell types are dispersed throughout the animal, including opsin-expressing cells (Layden et al., 2016; McCulloch et al., 2023).

Though *N. vectensis* is eyeless, larval dispersal, adult locomotor activity, and sexual reproduction are all light-dependent (Layden et al., 2016; Tarrant et al., 2019). *N. vectensis* also has among the highest number of opsins found in animals, at 29 (McCulloch et al., 2023). Additionally, *N. vectensis* have opsins from three groups: cnidopsins, which are sister to bilaterian xenopsins, the **A**nthozoan **S**pecific **O**psins **II** (ASO-II) which are sister to canonical visual c-opsins, and the **A**nthozoan **S**pecific **O**psins **I** (ASO-I) which are sister to all other animal opsins (Fig. 1B) (McCulloch et al., 2023). In contrast, medusozoans only have one opsin class, the cnidopsins, which are not sister to c- or r-opsins, making it a challenge to compare these pathways without direct orthology (Fig. 1B) (Macias-Munõz et al., 2019; Suga et al., 2008). The anthozoan-specific opsin diversity could also be reflected in distinct signaling pathways present in the variety of opsin-expressing cell types in *N. vectensis* (McCulloch et al., 2023). Understanding the extent of conservation in phototransduction cascade proteins in *N. vectensis* could help us trace the origins of the phototransduction cascade in animals.

**Figure 1.**
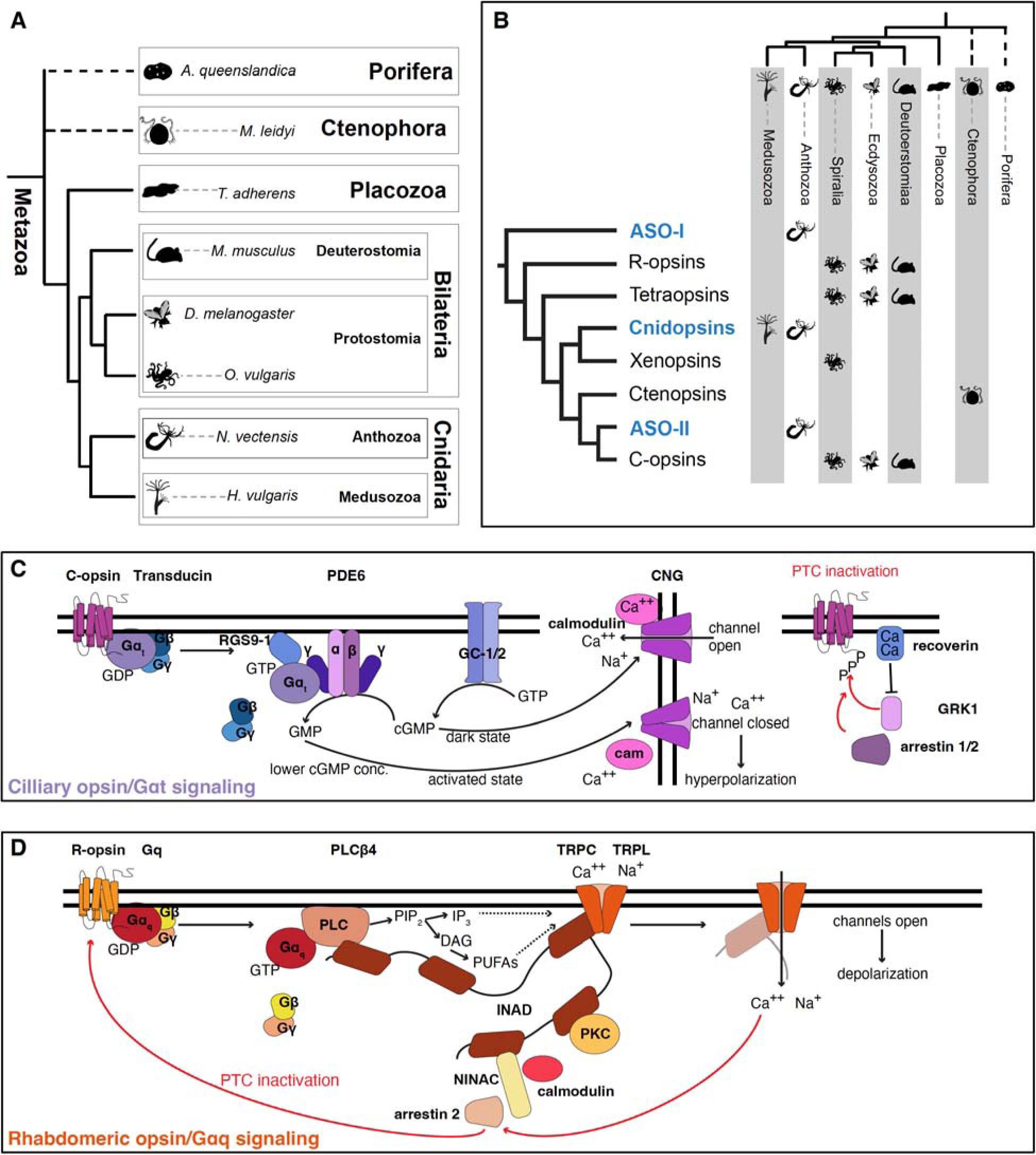
Major animal and opsin lineage relationships. **A)** Animal tree showing the major animal groups and the representative species from each group highlighted in this study. **B)** Current hypothesis of animal opsin relationships with presence or absence of each opsin group in each animal lineage indicated by the symbol. Cnidopsins and xenopsins together form a monophyletic clade, and ASO-II and c-opsins together form a monophyletic clade. Cnidarian-specific groups are highlighted in blue. Tree adapted from McCulloch et al. 2023. **C)** Schematic of the c-opsin phototransduction cascade and the proteins investigated in the current study. **D)** Schematic of the r-opsin phototransduction cascade and the proteins investigated in the current study.

Given that signals of at least partial conservation of phototransduction cascades are seen across animals, we hypothesize *N. vectensis* would make use of these same conserved pathways for phototransduction. The current study aims to identify the presence of orthologs of known phototransduction cascade genes in *N. vectensis* as a first step in characterizing the phototransduction cascade members in this understudied clade. We searched exhaustively for the major components of known phototransduction cascades across major animal groups with an emphasis on adding anthozoan representatives for each tree. We identify several lineage-specific duplications among these highly conserved genes, many unique to Anthozoa. This is the first attempt at characterizing phototransduction gene candidates in any anthozoan and an important step toward functionally characterizing these genes in the future.

## 2. Material and Methods

## 2.1 Sequence Retrieval

Accession numbers for *Hydra vulgaris* candidate phototransduction genes were obtained from Table 1 of (Macias-Munõz et al., 2019) and their coding sequences were retrieved from NCBI GenBank. To find the *N. vectensis* candidate proteins, BLASTx searches were performed using the *H. vulgaris* coding sequences as bait into the *N. vectensis* annotated proteins from the v2 genome in Geneious (Altschul et al., 1990; Zimmermann et al., 2020). Reciprocal BLAST searches were performed for each top query hit into the *Hydra vulgaris* v2 mRNA gene model. If the top query hit was not the same, reciprocal BLAST searches were repeated until the same result was obtained, and all results were added. For each search, the top hits with significant percent similarity (>60%) and query coverage (>50%) were saved.

**Table 1.** Presence of phototransduction cascade orthologs in representative species.

| Species | Opsin # | G alpha # | Effector enzyme # | Channel # |
| --- | --- | --- | --- | --- |
| c-opsin cascade | c-opsin | Gai/t | PDE6 | CNG |
| *M. musculus* | 4 | 2 | 3 | 6 |
| Insect | 1 | 1 | 1 | 4 |
| *N. vectensis* | 15 | 1 | 1 | 5 |
| *H. vulgaris* | 0 | 1 | 1 | 1 |
| r-opsin cascade | r-opsin | Gaq | PLC beta | TRPC |
| *M. musculus* | 1 | 1 | 1 | 7 |
| Insect | 7 | 1 | 2 | 3 |
| *N. vectensis* | 0 (2*) | 1 | 2 | 8 |
| *H. vulgaris* | 0 (0*) | 1 | 3 | 1 |

\* ASO-I opsins, which in some phylogenetic hypotheses are sister to r-opsins, though this relationship is still debated.

Following initial identification of *N. vectensis* candidate proteins, BLASTp searches were performed on NCBI using the *N. vectensis* sequences as bait. The sequences were first searched on the UniProtKB/Swiss-Prot (swissprot) data, followed by the Reference proteins (refseq_protein) and non-redundant protein sequences (nr) if insufficient query hits were shown. The *Nematostella vectensis* sequences were searched in the following taxa: *Mus musculus* (taxid:10088), *Danio rerio* (taxid:7955), *Drosophila melanogaster* (taxid:7227), *Xenopus tropicalis* (taxid:8364), *Mnemiopsis leidyi* (taxid:27923), *Trichoplax adherens* (taxid:10227), *Crassostrea virginica* (taxid:6565), *Limulus polyphemus* (taxid:6850), *Octopus bimaculoides* (taxid:37653), *Acropora millepora* (taxid:45264), *Exaiptasia pallida* (taxid:2652724), Cubozoa (taxid:6137), Scyphozoa (taxid:6142), Hydrozoa (taxid:6074), Zoantharia (taxid:6102), and Octocorallia (taxid:6132). The top 1-6 query hits for each taxon were manually selected for each species from each *N. vectensis* candidate gene BLAST result. Redundant sequences with the same accession numbers, identical protein sequences, and similar isoforms were removed. When the *N. vectensis* sequences did not yield sufficient BLAST results for a species, sequences found in other similar species were used as the bait. The *N. vectensis* candidate protein sequences were also searched in the genome assemblies of *Alatina alata* (Ohdera et al., 2019) and *Aurelia aurita* (Gold et al., 2019), and the top query hits were saved. Additional sequences were obtained from alignments from (Krishnan et al., 2015; Lagman et al., 2022; Moroz et al., 2023; Peng et al., 2015; Vöcking et al., 2022) and redundant sequences were not included.

For the G protein □ genes, *N. vectensis* sequences were retrieved from Interpro (Paysan-Lafosse et al., 2023) using the domain accession numbers PS50058 and PF00631. Adenylyl cyclase sequences were retrieved from (Vöcking et al., 2022) and used as bait to search on NCBI BLAST in the species listed previously.

### 2.2 Alignments and Phylogenetic Tree Construction

Amino acid sequences from the resulting hits were aligned with the *N. vectensis* candidate gene sequences on Geneious Prime with the MAFFT Alignment v7.450 using default parameters (Katoh and Standley, 2013).

Pseudo-maximum likelihood trees were generated from the MAFFT alignments using FastTree v2.1.11 (Price et al., 2010) with default settings. These trees were reviewed to identify the closest outgroups and to check that representative species were present in each clade within each tree. Sequences falling outside of outgroups, orphans, and highly divergent sequences were removed. When previously uncharacterized subclades within each gene tree were identified in other animals but not *N. vectensis,* additional BLAST searches were performed and added to the alignment. If newly added *N. vectensis* sequences were identified in the gene tree but not in any known subclades, additional BLAST searches in the above species were performed to confirm the presence or absence of orthologs in each animal lineage. Alignments and FastTrees were made iteratively as sequences were added and removed to finalize the trees.

Finalized alignments were realigned with MAFFT using the same settings as previously. For each gene family, a maximum likelihood tree was constructed using IQtree2 (Minh et al., 2020) on the Minnesota Supercomputing Institute’s Agate cluster using the following command: iqtree -s <ALIGNMENTNAME.phy> -st AA -nt AUTO -v -m TEST -bb 1000 -alrt 1000. This allowed for auto-choosing the best model of protein evolution for each tree, based on the lowest likelihood score from the Bayesian Information Criterion test. IQtree results yield both UltraFast bootstrap (UFbs) and aSH-LRT support values. Branches are considered highly supported with 80% aSH-LRT /95% UFbs support and were represented with a red circle on the Figures. Tree Figures were created using iTOL and inkscape (Letunic and Bork, 2021). Species icons for Placozoa and Porifera were obtained from phylopic.

### 2.3 *In situ* Hybridization

The G protein α subunit VI probe was generated from a PCR product on *N. vectensis* cDNA, using the primer pair: Fwd:ATTCAGGCAAAAGCACGTTT Rev:GGGATGCGAAAAATACCACC. The PCR product was ligated to pGEM-T Easy vector backbone (Promega) and DIG-labeled RNA probes were synthesized using T7 megascript kit (Ambion) and cleaned up using a Zymo RNA cleanup kit, according to previously published methods (McCulloch et al., 2023). *In situ* hybridization was performed using a standard *N. vectensis* protocol as previously published (McCulloch 2023). Animals were mounted on a microscope slide and chromogenic staining was visualized with a Nikon90i using DIC optics.

### 2.4 mRNA Expression Patterns

*N. vectensis* phototransduction candidate proteins that were confirmed phylogenetically were then identified on the *N. vectensis* embryonic and regeneration transcriptome database, NvERTx.4 (Warner et al., 2018) using tBLASTn default parameters. The result with 100% match was selected and the regeneration expression data was retrieved. Developmental expression data was retrieved from previously published transcriptomic data (McCulloch et al., 2023). For details on the assembly and mapping methods please refer to the original references ((McCulloch et al., 2023; Warner et al., 2018) The regeneration data is in normalized “counts,” available on NvERTx, while the developmental time series is shown in normalized TPM.

## 3. Results and Discussion

Opsins evolved early in animals and are present in nearly all major groups (Fig. 1A,B). Canonical ciliary and rhabdomeric phototransduction cascades make use of distinct opsins and other proteins in the pathway (Fig. 1C,D). Although c- and r- pathways are best known in vertebrates and insects respectively, both animal lineages have components of both pathways (Table 1). A summary of *N. vectensis* and *H. vulgaris* c- and r- pathway members show both cnidarian species also have the core components of these pathways (Table 1). Expression levels of all the transcripts identified in this study during regeneration and embryonic development are summarized in heatmaps (Fig. 2).

**Figure 2.**
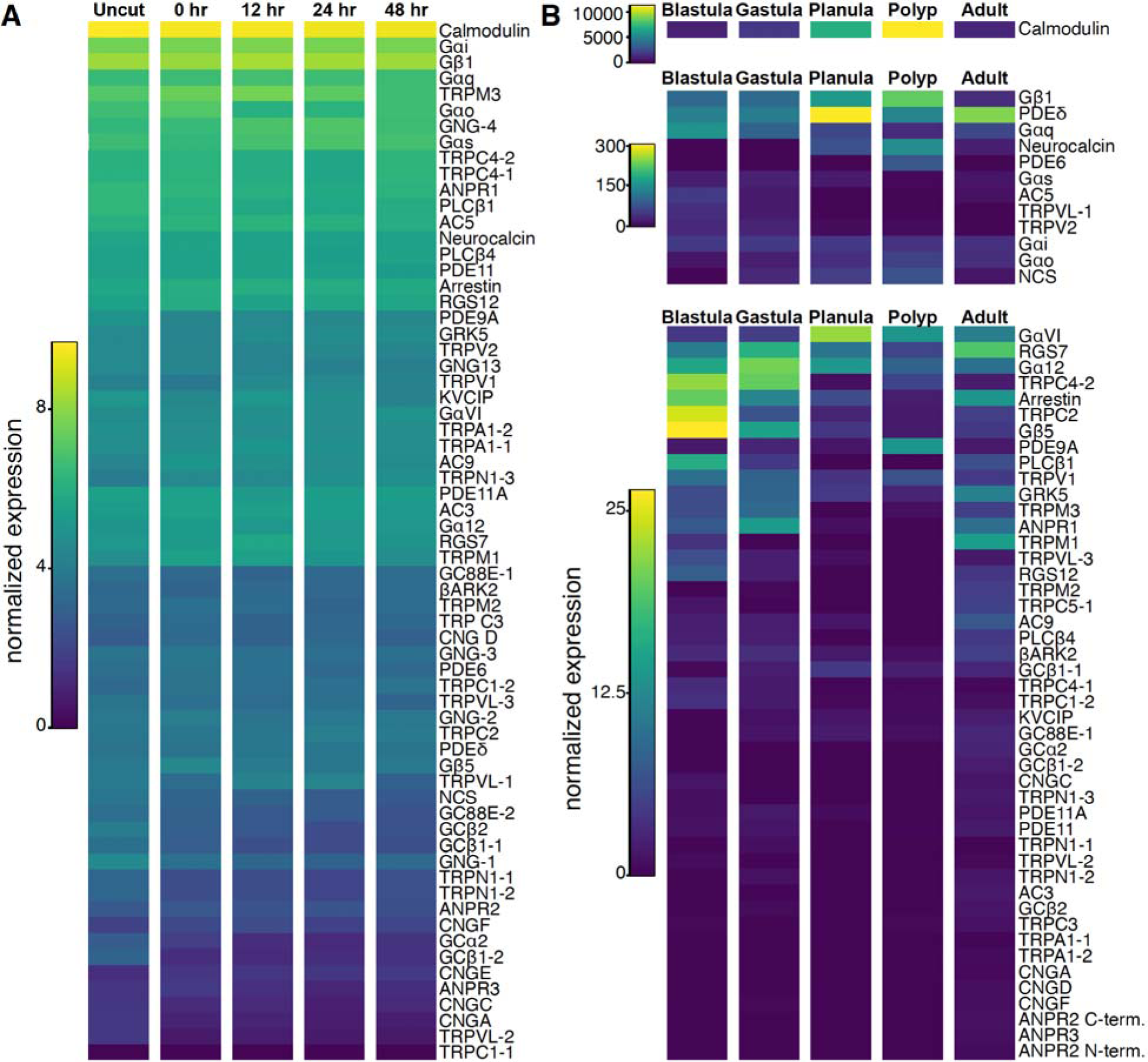
mRNA expression levels of phototransduction genes in regeneration and development. **A)** Expression levels from timecourse series at different stages of whole regenerating oral pole from NvERTx (Warner et al. 2015). **B)** Expression levels from timecourse series at different stages of development from whole animals, from blastula to adult. Data is from (McCulloch et al. 2023). Different scales are used for the top expressing genes for visual clarity.

### 3.1 Initiation of the phototransduction cascade

All phototransduction cascades across animals are initiated by a rhodopsin absorbing light and activating the associated heterotrimeric G protein, constituted of an α, β, and □ subunit. Upon photon absorbance, the Gα separates from the β and □ subunits, and functions as a GTPase to activate an effector enzyme, which can differ depending on the type of phototransduction cascade (Shichida and Matsuyama, 2009).

#### 3.1.1 Heterotrimeric G protein

The Gα subunit is the most studied of the G protein complex. Best known for activating cGMP phosphodiesterase (PDE) in ciliary opsin cells and for activating phospholipase C (PLC) in rhabdomeric opsin cells. While the β and □ subunits participate in regulation of the cascade, much less is known about their function in any phototransduction cascade outside of vertebrates.

##### Gα subunit

There are five known major groups of Gα subunits in animals, of which three are known to be involved in phototransduction (Lokits et al., 2018). Within these major groups, duplications and subsequent lineage-specific Gα subunits have evolved, such as the vertebrate Gαt, which is a duplicate of the ancestral Gαi group (Lagman et al., 2012). Well-studied pathways involve the Gαq protein which activates the r-opsin signaling cascade, and the Gαi/Gαt family found in retinal rods and cones (Shichida and Matsuyama, 2009).

We surprisingly found that *N. vectensis* has 6 Gα orthologs, five in the families Gαs, Gαo, Gαq, Gαi, Gα12, and one in a highly supported novel clade within the Gα tree (Fig. 3A, Fig. S1). Our tree places this group sister to the Gαq and Gα12 clade, although this relationship has low support. Domain analysis reveals that this unspecified *N. vectensis* gene retains the diphosphate binding (P-) loop (GXGESGKS), Mg^2+^ binding domain (RXXTXGI and DXXG), and guanine ring-binding motifs (NKXD and TCAT) within the GTPase domain characteristic of Gα protein structure (Oldham and Hamm, 2006). An ortholog of this gene was previously identified in the coral *Acropora palmata,* though no phylogenetic classification was attempted (Mason et al., 2012). Our novel gene is highly conserved with this coral Gα. We identify additional anthozoan representatives in this new clade, but unexpectedly we also found multiple spiralian sequences and a single ortholog in *Branchiostoma belcheri* (Fig. 3A, Fig. S1). However no results were found for any ecdysozoan, any other deuterostome, or any medusozoan. Because the gene is present in both Cnidaria and Bilateria, this novel clade was likely present in their common ancestor and subsequently lost in other animal lineages. We propose to refer to this sixth group as GαVI.

**Figure 3.**
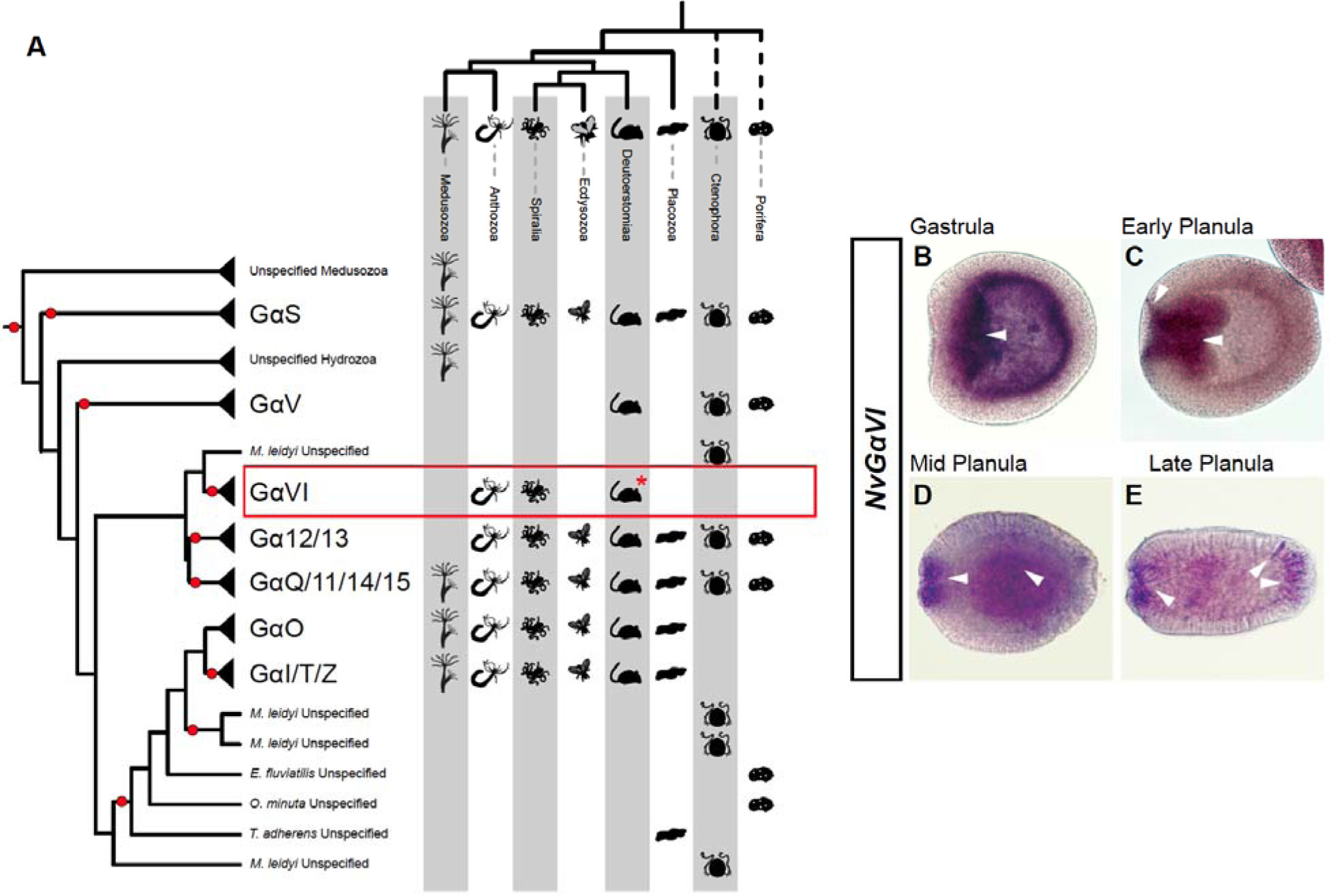
Gα subunit protein evolution in animals. **A)** Maximum-likelihood G_α_ subunit tree with major subgroups collapsed and individual branches from undefined sponge and ctenophore sequences. Presence of each subfamily in each animal lineage is indicated by animal symbol. A novel group, GαVI, is indicated by a red box. Asterisk, subunit is found in Deuterstomes, but only found in *Branchiostoma*. Red circles indicate highly supported nodes with 80% aSH-LRT/95% UFbs support. **B-E)** Spatial expression of the novel *NvGaVI* mRNA in *N. vectensis* development. **B)** At gastrula stage, expression is highest in the pre-endodermal plate (arrowhead) which is invaginating and is beginning to form the endoderm. **C)** By early planula stage expression is around the lip of the mouth, the elongating pharynx ectoderm, and the expanding endoderm (arrowheads). **D)** By mid planula expression is seen in the cells surrounding the mouth and broadly in the interior endoderm (arrowheads). **E)** By late planula expression is still found around the mouth, and in the sensory apical organ (arrowheads). Some expression is still seen in broadly in the endoderm. In all panels, mouth is to the left, apical organ/aboral to the right.

To identify expression patterns of this novel gene in *N. vectensis* development, *in situ* hybridizations were performed (Fig. 3B-F). We find that early in gastrulation, *NvG*α*VI* expression is concentrated in the pre-endodermal plate, and continues throughout gastrulation as this invaginates and the endoderm is formed (Fig. 3C-D). Later, in the larval planula stages, *NvG*α*VI* is expressed broadly in the endoderm and specifically in the ectodermal lip where the cells begin to internalize and form the pharynx (Fig. 3E-F). Expression is also seen in later planula stages in the sensory apical organ (arrowheads, Fig. 3F). No expression could be seen in later stages although RNA-seq data suggest some expression in developing and regenerating polyps (Fig. 2). Some opsins are known to be expressed near the mouth and in the apical organ at these stages, including *NvASOII-8b* and *NvASOI-2* (McCulloch 2023), however it is still unknown whether there is a functional link between this novel GαVI subunit and opsins.

G proteins are involved in many types of cellular signaling so we cannot rule out that these orthologs may be used for functions other than photoreception. However evidence of mulltiple Gα family members in cnidarian phototransduction suggests these might function in *N. vectensis* phototransduction as well. The Gαs protein of the box jelly *Tripedalia cystophora* may regulate cAMP concentrations in response to light (Koyanagi et al., 2008). In the coral *A. palmata*, two ASO-II opsins were capable of activating the *A. palmata* Gαq ortholog, while one of these could activate the unspecified Gα protein that we now classify as the novel GαVI (Mason et al., 2012). *N. vectensis* has 15 ASO-II paralogs and 12 cnidopsin paralogs, so it is possible some of these could use Gαs, Gαq, or GαVI for phototransduction.

##### Gβ subunit

Five major Gβ subunits are known in animals, including Gβ1 expressed in rod photoreceptors, Gβ3 (β-transducin) expressed in cone photoreceptors (Dexter et al., 2018; Peng et al., 1992). Our tree aligns with previous research showing that Gβ homologs are split into two major clades in animals: Gβ5, and Gβ1 which has been duplicated multiple times in vertebrates (Gβ1-4) (Fig. S2). Similar to protostomes, we found that *N. vectensis* has one homolog in each of these major clades (Fig. S2). In contrast to *N. vectensis*, *H. vulgaris* has three closely related copies of the gene within the Gβ1-4 group which likely arose from species-specific duplications of the *H. vulgaris* Gβ1 gene.

##### G□ subunit

At least 12 G□ subunit genes have been identified in vertebrates. The GNG cluster contains 2 deuterostome-specific subgroups with 8 paralogs among vertebrates, and a single ortholog in protostomes (Krishnan et al., 2015). One GNG13 ortholog is typically present in all animals except sponges and ctenophores, while the G□ transducin (GNGT) family is vertebrate specific. Previous work identified 2 *N. vectensis* genes in the GNG cluster and 1 GNG13 homolog (Krishnan et al., 2015).

Using domain searches in Pfam rather than BLAST, we confirmed that *N. vectensis* has the 3 G□ genes previously identified and identified 2 additional G□ sequences (Fig. 4, Fig. S3). These fall into two subgroups outside of the previously identified G□ subclades. One sequence is in a cnidarian-specific group where the three *Hydra* duplicates are found, and another is found in an anthozoan-specific group. Whether the novel G□ subgroups are sister to the established groups is not highly supported, however the topology placing all of these in a single clade is highly supported (Fig. 4).

**Figure 4.**
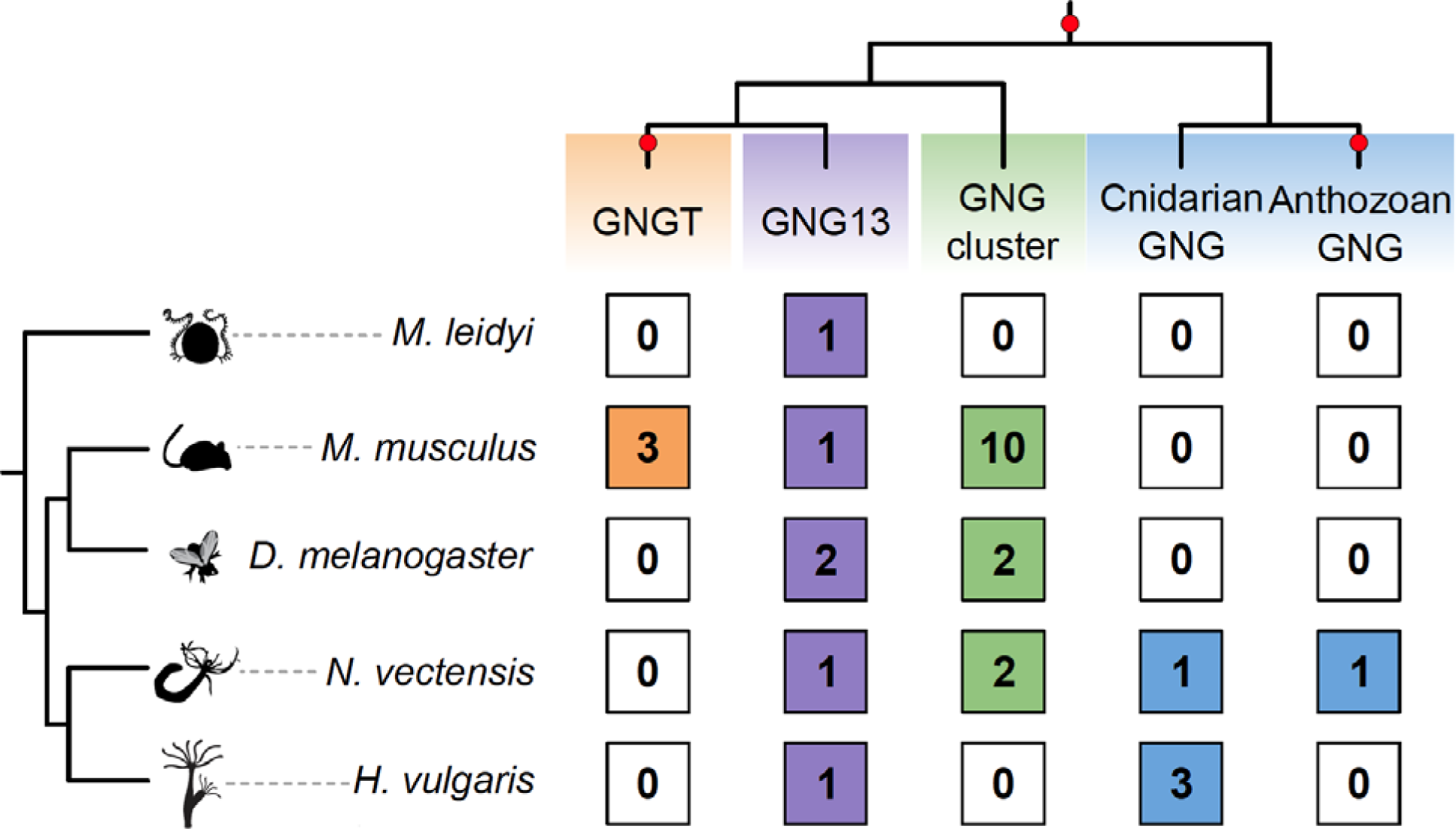
G_subunit protein family. Simplified maximum-likelihood G□ tree with major subgroups collapsed, top. The number of subfamily paralogs in each animal lineage is indicated by numbers in colored boxes. Red circles indicate highly supported branches with 80% aSH-LRT/95% UFbs support.

One of the reasons for this uncertainty is the short sequence length of the G□ gene (49-75 amino acids long for the *N. vectensis* genes), which makes phylogenetic inference a challenge. To confirm that the unspecified *N. vectensis* G□ genes might still have G protein function, we analyzed the *N. vectensis* sequences using InterPro database functional domain scans in Geneious. This analysis predicted intact G□ functional domains, confirming these could likely function in the heterotrimeric complex with other G protein α and β subunits. While the relationships between these different G□ genes is not certain, we show that *N. vectensis* has more G□ subunits than previously identified.

Previous research has shown both interchangeability and specificity in □ subunits with evidence that G□ subunit identity is not important in some signaling contexts *in vivo*, while in other cases a particular G□ subunit is required for a specific function (Dexter et al. 2018). It is thus possible the multiple *N. vectensis* G□ genes may be similar, both functioning in specific Gβү complexes for particular phototransduction cascades and also others that are interchangeable in other G protein signaling contexts.

#### 3.1.2 Effector enzymes

After activation by the Gα subunit, effector enzymes such as cGMP-PDE and PLC affect the concentrations of second messengers. Other effector enzymes include guanylyl cyclase (GC) in Gαo-opsin signaling in the ciliary photoreceptors of scallops (Del Pilar Gomez and Nasi, 2000) and adenylyl cyclase (AC) in chicken Gαi signaling via neuropsins (tetraopsins) in non-typical photoreceptor cells of the retina and brain (Yamashita et al., 2010). AC has also been implicated in Gαs signaling in box jelly phototransduction (Koyanagi et al., 2008).

##### Phospholipase C

In *Drosophila,* PLCβ4 hydrolyzes phospholipid phosphatidylinositol 4,5-bisphosphate (PIP2) to inositol 1,4,5-triphosphate (InsP3) and diacylglycerol (DAG) upon activation by Gαq (Hardie and Raghu, 2001). These molecules both act as second messengers to change the concentrations of calcium and other metabolites within the cell, leading to the opening of transient receptor potential channels (TRPs). We show that *N. vectensis* has the same two orthologs as the *Drosophila* PLCβ genes (Fig. S4). The presence of both a Gαq gene and PLCβ4 homolog shows that *N. vectensis* have the necessary components to utilize a signaling pathway similar to that of the rhabdomeric pathway.

##### Cyclic GMP-Phosphodiesterase (cGMP-PDE)

In vertebrate retinal ciliary photoreceptors, the active cGMP-PDE6 complex increases levels of the second messenger cGMP, which then closes cyclic nucleotide gated (CNG) channels in the plasma membrane. The vertebrate PDE6 is made up of one α subunit, one catalytic β subunit, and two inhibitory □ subunits. The group sister to the vertebrate PDE6 subunits contains the single *Drosophila* cGMP-specific PDE6 ortholog. Cnidarians appear to also have an ortholog to PDE6, but this is unclear due to low support (Fig. S5). If PDE6 signaling is a vertebrate-specific innovation, *N. vectensis* could be more similar to invertebrate c-opsin signaling, with ASO-II opsins signaling through GC rather than PDE6.

##### Guanylyl Cyclase

The guanylyl cyclase (GC) enzyme family is split into transmembrane and soluble subfamilies. The transmembrane GCs, also called atrial natriuretic peptide receptors (ANPR), can activate or inhibit phototransduction via calcium- and cGMP-dependent feedback loops (Dizhoor et al., 1994; Koch and Stryer, 1988). The transmembrane GCs in vertebrate phototransduction are part of a vertebrate-specific subfamily called retinal GCs while evidence from scallop suggests an ANPR subfamily member is involved in phototransduction (Del Pilar Gomez and Nasi, 2000; Potter, 2011). We found that *N. vectensis* has 3 transmembrane GC genes and 6 soluble GC genes (Fig. 5A, Fig. S6). One of these NvGC paralogs is in a cnidarian-specific group sister to the bilaterian ANPR family, while the other two are found in two additional cnidarian- and anthozoan-specific groups within the transmembrane GCs (Fig. 5B). We hypothesize that the NvANPR could function in a conserved role in phototransduction like the scallop ANPR, potentially in NvASO-II (c-opsin-related) expressing cells. The functions of the cnidarian transmembrane GCs are unknown. However gene expansions in the GC and other phototransduction-related gene families could correlate with the expanded set of *N. vectensis* opsins and be used for different phototransduction contexts.

**Figure 5.**
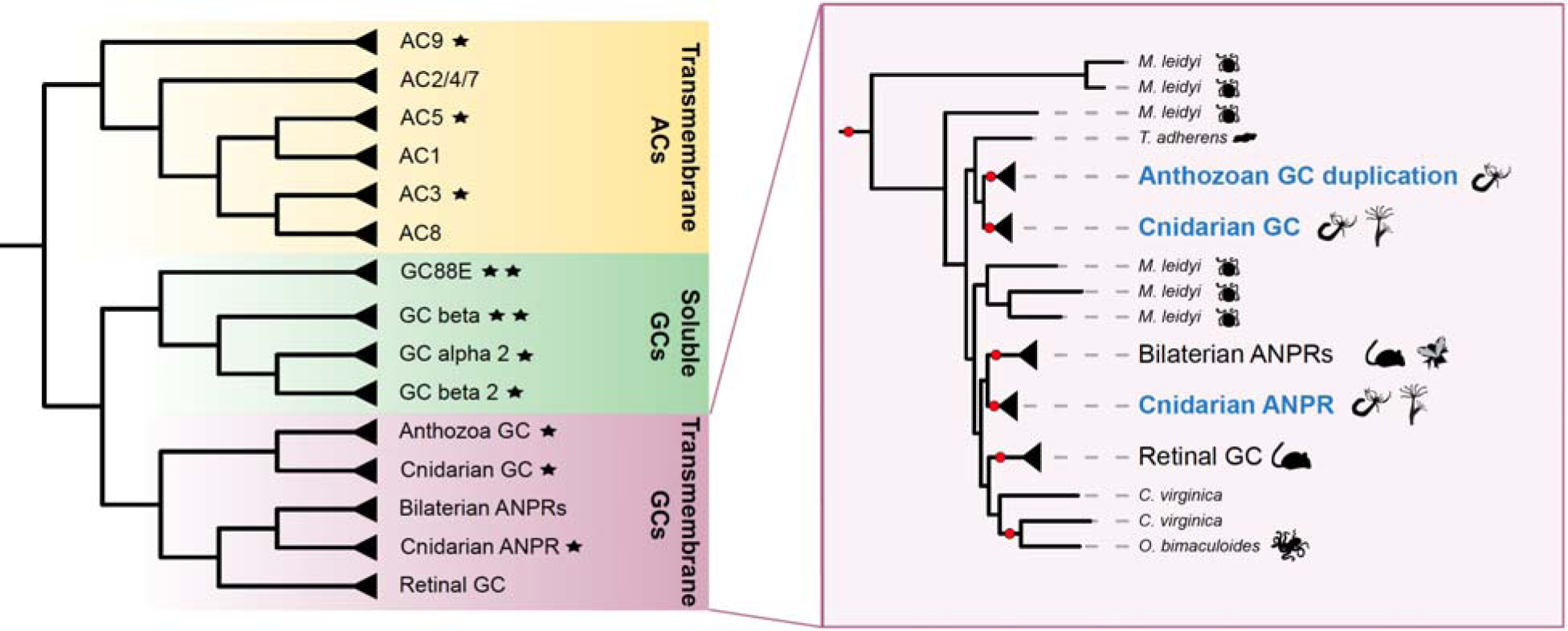
Anthozoan expansions in the AC/GC/ANPR protein family. **A)** Maximum-likelihood tree with major AC/GC/ANPR subgroups collapsed and individual branches with no clear group from Ctenophora, Placozoa, and Spiralia removed. **B)** Transmembrane GCs in the tree expanded, showing many orphans with no clear group from Ctenophora, Placozoan, and Spiralia. Groups in blue show cnidarian and anthozoan-specific groups. Red circles indicate highly supported branches with 80% aSH-LRT/95% UFbs support.

##### Adenylyl cyclase

While cGMP is the most studied secondary messenger in vertebrate c-opsin signaling, cAMP signaling via AC5/6 has been suggested as having a major role in the ancestral chordate phototransduction cascade (Lamb and Hunt, 2017). Recent evidence points to this with a role of cAMP in vertebrate r-opsin signaling. In mice, AC2 is activated by Gaq in the melanopsin pathway (Chen et al., 2023). In the box jelly *T. cystophora*, a cnidopsin-Gαs pathway activates an AC enzyme that affects cAMP concentrations (Koyanagi et al., 2008). It is currently unknown which specific AC type is involved, however AC3 has been suggested due to the similarities between the jellyfish phototransduction cascade and the vertebrate olfactory signaling cascade (Firestein, 2001).

We found that *N. vectensis* has homologs of AC types 5/6, 3, and 9 (Fig. 5A). This confirms previous research indicating cnidarians only have these three AC types (Vöcking et al. 2022). AC3 and AC5/6 are likely candidates in the phototransduction cascade in *N. vectensis,* possibly through cnidopsin signaling. Overall, patterns of GC and AC participation in various phototransduction cascades suggest that these enzymes can be repeatedly recruited and swapped for one another over the course of animal evolution. Their prevalence across several types of phototransduction cascade suggests they were recruited early while the vertebrate c-opsin cascade relying on PDE6 may be a later innovation.

#### 3.1.3 Ion Channels

When effector enzymes alter the concentrations of second messengers, ion channels open or close in response, leading to voltage changes and propagation of the light signal. In c-opsin expressing ciliary photoreceptors, these are CNG channels while r-opsin expressing rhabdomeric photoreceptors utilize TRP channels.

##### Cyclic nucleotide gated (CNG) channels

CNG channels are thought to be the ancestral ion channel used in phototransduction before the radiation of metazoans (Lagman et al., 2022). Dependent on the type of signaling, CNG channels are either opened or closed in response to cascade activation (Fain et al., 2010). Generally, three α subunits and one β subunit are used to assemble the channel, and these can be mixed and matched (Lagman et al., 2022). There are six subunits of CNG channels in animals, and anthozoans have the most of any animal group with five (Lagman et al. 2022). We confirm that one subunit, CNGE, is anthozoan-specific (Fig. 6, Fig. S7). CNG channels are likely involved in *N. vectensis* phototransduction, as *Hydra magnipapillata* CNG channels are implicated in the contractile photo-response (Plachetzki et al., 2010).

**Figure 6.**
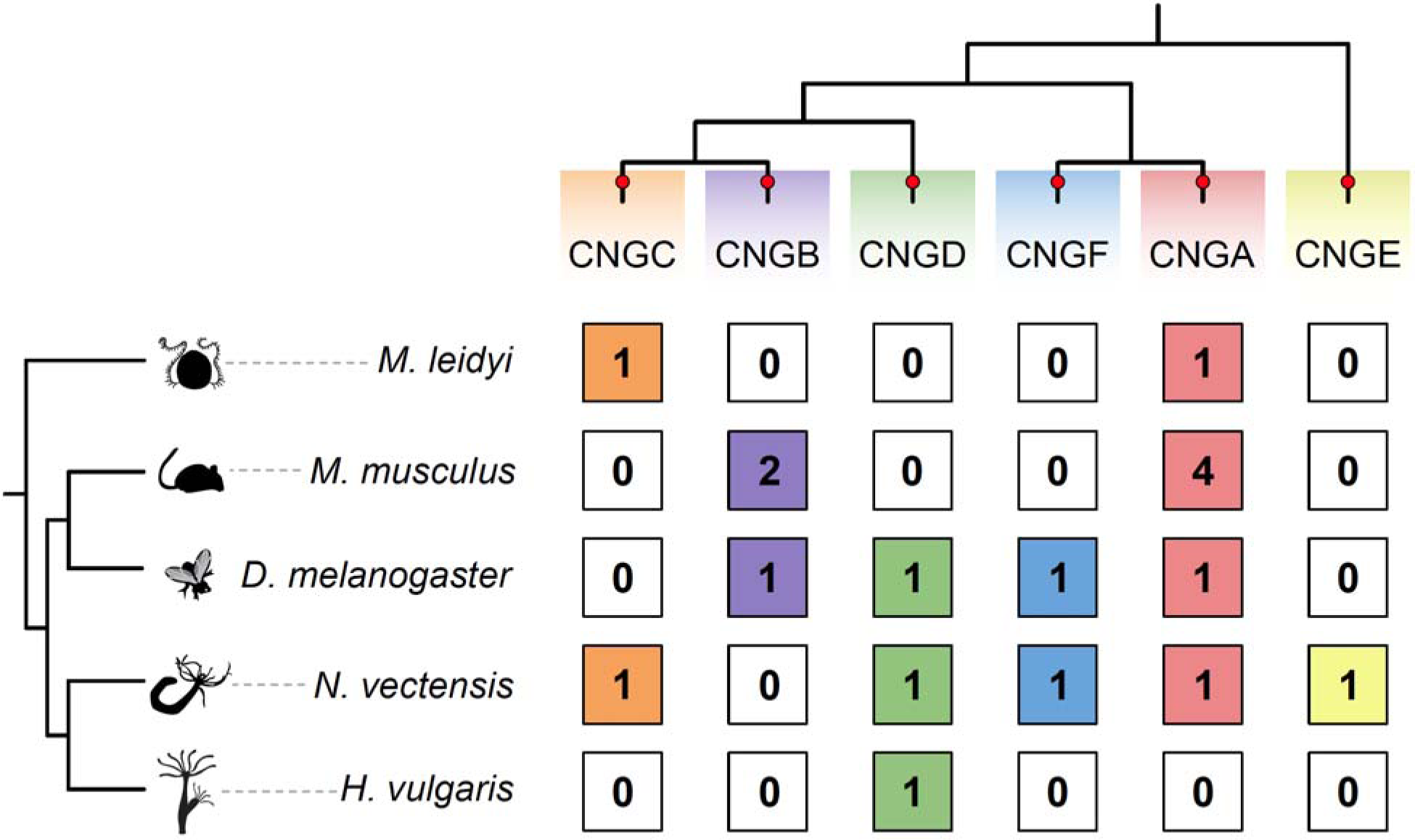
CNG subunit protein family. Simplified maximum-likelihood CNG tree with major subgroups collapsed, top. The number of subfamily paralogs in each animal lineage is indicated by numbers in colored boxes. Red circles indicate highly supported branches with 80% aSH-LRT/95% UFbs support.

Given these subunits form heterotetrameric channels in other animals, the large number of subunits found in Anthozoa could allow for high combinatorial diversity of CNG channels and their functional contexts. Our bulk RNA-seq data shows that CNG subunits are among the lower expressing transcripts in both development and regeneration (Fig. 2). However *N. vectensis* single cell transcriptomic data shows multiple subunits were co-expressed in the same cell types in nearly all tissue types, including neural and glandular cell types (Lagman et al., 2022; Sebé-Pedrós et al., 2018). Together this suggests CNG subunits are not highly expressed in bulk RNA-seq data because they are expressed in relatively fewer specialized cell types, and distinct combinations could be specifically expressed in sensory and photoreceptor cell types.

##### Transient receptor potential channels

TRPC channels are the main type of TRP channel in *D. melanogaster* r-opsin phototransduction, which open in response to cascade activation (Montell, 2005). TRP channels are known to be involved in a variety of sensory contexts in animals and may have been co-opted specifically into the r-opsin/Gq phototransduction cascade more recently in protostomes (Plachetzki et al., 2010), although without more functional studies in Cnidaria, especially Anthozoa, we cannot rule out a loss of Gq/TRP signaling in vertebrates.

We focused on TRPC channels which are the only TRPs known to be involved in phototransduction. Our phylogenetic analysis revealed a large anthozoan-specific expansion in TRPC channel homologs. *N. vectensis* has a total of 8 TRPC genes, while *H. vulgaris* has one TRPC gene (Fig. 7A, Fig. S8) (Peng et al., 2015; Vöcking et al., 2022). We find that previously identified cnidarian TRPCs are split into two groups, one of which is pan-cnidarian, while the other is anthozoan-specific (Fig. 7B, green and orange). The three *N. vectensis* TRPC genes previously found and one additional novel paralog are all sister to bilaterian TRPC. We also identified another anthozoan-specific TRPC group that falls sister to the previously defined groups and has four additional *N. vectensis* sequences (Fig. 7B, blue). We show that *N. vectensis* has multiple TRPC orthologs as part of either an anthozoan-specific expansion and divergence or retention of ancient paralogs of this ion channel subfamily. *N. vectensis* could use their many CNG and TRP channels in distinct signaling mechanisms, including potentially a variety of distinct phototransduction contexts.

**Figure 7.**
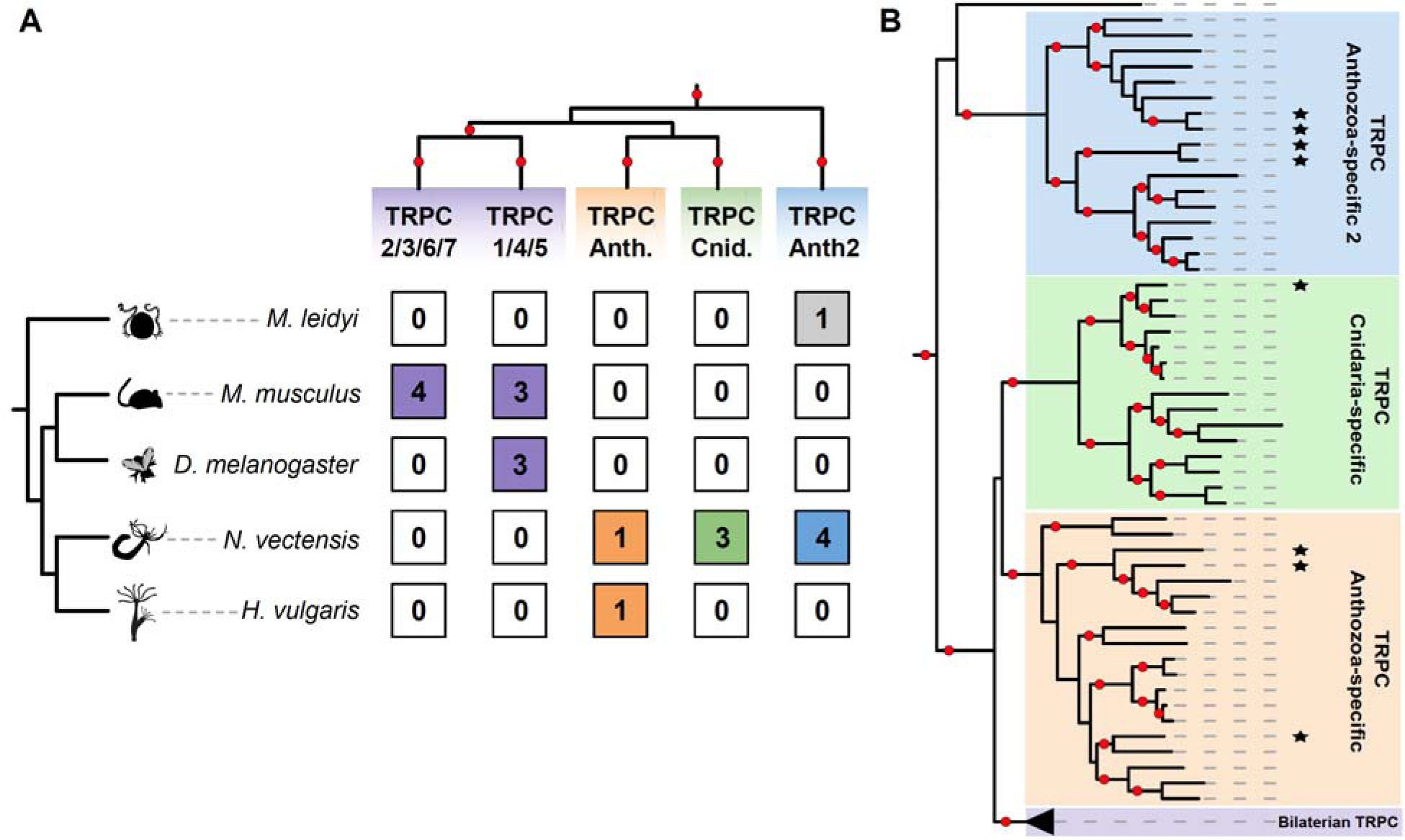
TRP channel evolution Anthozoa. **A)** Simplified maximum-likelihood TRPC tree with major subgroups collapsed, top. The number of subfamily paralogs in each animal lineage is indicated by numbers in colored boxes. Gray box is for the ctenophore sequence which is not supported in any clade but is sister to the Anthozoa 2 group. **B)** Expanded maximum-likelihood tree of TRPC subfamilies focused on cnidarian-specific expansions. Stars next to branches indicate the *N. vectensis* sequences. Red circles indicate highly supported branches with 80% aSH-LRT/95% UFbs support.

### 3.2 Regulation of the phototransduction cascade

#### G protein-coupled receptor kinases

The superfamily of G protein-coupled receptor kinases (GRKs) phosphorylate a variety of GPCRs and includes the visual GRKs rhodopsin kinase (GRK1) and GRK7. GRK1 is vertebrate-specific and has a role in recovery of the vertebrate ciliary phototransduction cascade by phosphorylating rhodopsin and allowing the inhibitory arrestin protein to bind (Lyubarsky et al., 2000). GPRK1 in *Drosophila* has a function analogous to vertebrate rhodopsin kinase (Lee et al., 2004). However DmGPRK1 is phylogenetically placed with the β-adrenergic receptor kinase group GRK2/3.

Another GRK, bovine GRK5 is expressed in the retina and can phosphorylate rhodopsin in a light-dependent manner when expressed in insect cells (Premont et al., 1994). We found that *N. vectensis* has homologs for both GRK5 and DmGPRK1, either of which could be involved in regulating G protein-coupled receptor activity during the phototransduction cascade. We confirm that GRK1 is vertebrate specific and that DmGPRK1 is included with the β adrenergic receptor kinases GRK2/3 clade (Fig. S9).

#### Arrestin

Visual arrestin is an essential protein in the termination of visual signaling that inactivates GPCRs and halts rhodopsin activity across Metazoa. Vertebrates have four arrestin proteins, two of which function in phototransduction termination in rods and cones (Lamb et al., 2018; Wilden et al., 1986). Protostomes have two arrestins with no 1:1 orthology with vertebrate arrestins, one of which is involved in phototransduction (Gurevich and Gurevich, 2006; Swardfager and Mitchell, 2007). In contrast, we identified only one arrestin gene in *N. vectensis* and other cnidarians (Fig. S10). This cnidarian arrestin has been shown to function similarly to bilaterian arrestins and plays a role in terminating G protein signaling in phototransduction (Plachetzki et al., 2012). The presence of an arrestin protein with similar function in both Bilateria and Cnidaria suggests that it has a highly conserved role in the termination of phototransduction in animals, including *N. vectensis*.

#### Calcium binding proteins

Calcium ions have a variety of roles in G protein signaling, including in multiple phototransduction cascades (Fain et al., 2010). Calcium-binding proteins regulate calcium ion concentration and distribution, playing important roles in the regulation of phototransduction (Ikura, 1996; Tanaka et al., 1995). The neuronal calcium sensor proteins are phylogenetically and functionally related (Burgoyne and Weiss, 2001). Calmodulin is a calcium binding protein that interacts with both CNG and TRP channels mediating positive and negative feedback mechanisms (Babu et al., 1988; Bej and Ames, 2022; Sun et al., 2018; Trudeau and Zagotta, 2004). In vertebrates neurocalcin regulates the GC enzyme (Dizhoor et al., 1994) and neurocalcin may also inhibits rhodopsin in fly eyes in a manner similar to recoverin in vertebrates (Faurobert et al., 1996). We found that *N. vectensis* has orthologs for calmodulin and neurocalcin, and two additional unspecified orthologs in the calcium binding family (Fig. S11). Calmodulin is also the highest expressing transcript in both regeneration and development (Fig. 2). It is possible that calmodulin and neurocalcin act in broad contexts for calcium signaling in *N. vectensis,* including in phototransduction.

#### Regulator of G protein signaling 9

Regulator of G protein signaling (RGS) members of the R7 family (RGS6, RGS7, RGS9, RGS11) form complexes with other proteins and act to regulate the phototransduction response (Makino et al., 1999; Squires et al., 2018). We show that *N. vectensis* has RGS7 and RGS12 orthologs (Fig. S12). Pfam analysis reveals that the *N. vectensis* RGS7 maintains the functional domains characteristic of the R7 family, which could allow it to regulate N. vectensis phototransduction in a similar manner (Levay et al., 1999; Squires et al., 2018).

### 3.3 Expression of phototransduction genes in N. vectensis

Once we identified the phototransduction gene candidates in *N. vectensis*, we wanted to know how they were expressed over both developmental and regeneration time (McCulloch et al., 2023; Warner et al., 2018). Throughout development, calmodulin and the G protein α and β subunits were among the highest expressing genes, including the novel GaVI (Fig. 7A). Many TRP channels were also highly expressed, particularly the TRPV, TRPVL, and some TRPC genes, qualitatively much more so than the CNG channels. This could indicate that TRP channels have a broader role in multiple signaling contexts in the animal while CNG channels might be more specifically expressed in fewer cell types. The cGMP-PDE δ subunit was also very highly expressed, particularly during the planula stage of development potentially correlated with sensory signaling specific to this swimming life stage, as some opsins are. Many genes were also expressed during regeneration but none in obvious regeneration-specific patterns (Fig. 7B).

### 3.4 Discussion of N. vectensis phototransduction and evolutionary implications

Our results reveal that similar components required for phototransduction in the major c-opsin and r-opsin phototransduction cascades are present in the eyeless Anthozoa. We summarize our results in Table 2 by showing the number of paralogs for each gene family in select species. This aligns with the limited evidence from other cnidarians suggesting conserved phototransduction components (Koyanagi et al., 2008; Kozmik et al., 2008; Vöcking et al., 2022). Of note, *N. vectensis* consistently has more paralogs for most gene families relative to *Hydra*, and this correlates with *N. vectensis* having more opsin receptor classes (3) than *Hydra* (1). It is expected that vertebrates have more paralogs due to whole-genome duplications, however *N. vectensis* has more than even vertebrates for some gene families (TRPC, GC) and more than *D. melanogaster* for several gene families (TRPC, GC, ANPR, CNG, PDE, G_). This suggests that patterns of gene duplication leading to sub- and neofunctionalization are not exclusive to Bilateria and occurred in parallel in Cnidaria. Further investigation in overlooked lineages such as Anthozoa will continue to reveal previously unappreciated evolutionary history and the origins of complexity among animals.

**Table 2.** Summary of phototransduction homologs in representative species.

| Gene | *M. leidy* | *H. vulgaris* | *N. vectensis* | *M. musculus* | *D. melanogaster* |
| --- | --- | --- | --- | --- | --- |
| TRPC | 1 | 1 | 8 | 7 | 3 |
| TRPM | 0 | 6 | 3 | 8 | 1 |
| GC | 2 | 2 | 6 | 3 | 5 |
| AC | 2 | 4 | 3 | 8 | 12 |
| ANPR | 1 | 2 | 3 | 3 | 0 |
| Arrestin | 1 | 1 | 1 | 4 | 3 |
| CNG | 2 | 1 | 5 | 6 | 4 |
| GMP-PDE | 3 | 5 | 5 | 6 | 4 |
| G alpha | 11 | 4 | 6 | 17 | 8 |
| G beta | 3 | 2 | 2 | 5 | 2 |
| G gamma | 1 | 4 | 5 | 14 | 3 |
| PLC | 1 | 3 | 2 | 4 | 2 |
| GRKs | 2 | 2 | 2 | 4 | 2 |
| RGS (R-7 family) | 1 | 2 | 2 | 5 | 3 |

A novel Gα subunit, GαVI, was also identified in Anthozoa and Spiralia (and *Branchiostoma*) and is absent in the major vertebrate and ecdysozoan animal models. This suggests that losses have occurred in these animal lineages, and a lack of attention in cnidarian and spiralian groups left this subfamily previously unidentified. G proteins and their associated α subunits are involved with a diverse range of signaling mechanisms, so this novel GαVI may have important roles in these species including in phototransduction.

In Anthozoa, Medusozoa, and Bilateria, we see many of the same core phototransduction pathway components (opsins, Gα, effector enzymes, and channels). Some specific pathways, such as vertebrate c-opsin/Gαt signaling, have specialized by swapping paralogs or even adding novel enzymes to the cascade. When compared alone, it is impossible to say whether the highly specialized Gαq/r-opsin pathway and the Gαt/c-opsin pathway in insect and vertebrate eyes are *de novo* evolved or share ancestry. Our findings along with evidence that opsins diverged before the split of Bilateria and Cnidaria suggest that the phototransduction cascades across Metazoa could have arisen in their earliest common ancestor. This is further supported by similarities between medusozoan phototransduction and the vertebrate olfactory transduction cascade (Firestein, 2001; Koyanagi et al., 2008; Ramirez et al., 2016). Our data are in line with the hypothesis that the last common ancestor of animals had a general GPCR signaling pathway before opsins and olfactory receptors split, which then diversified as new sensory modalities evolved. With more sampling in new lineages and contexts, and with further functional data in the future, we are closer to understanding the origin and continued evolution of phototransduction in animals.

## 4. Conclusions: Implications for the phototransduction cascade in *N. vectensis*

Current evidence suggests both opsins and their respective phototransduction cascades evolved before the split of Bilateria and Cnidaria. By identifying well-known phototransduction cascade members in *N. vectensis,* we show the first evidence that Anthozoa possess the capacity to signal through similar cascades as Bilateria. Furthermore, by investigating a relatively understudied lineage, we have identified novel subgroups in otherwise well-studied gene families, such as GαVI and multiple novel G□ subunits, GC genes, and TRP channels in *N. vectensis* and Anthozoa. More functional work must be done to confirm these genes work together in a cascade. Even so, our findings highlight that broader taxonomic sampling in otherwise well-known gene families can uncover previously unknown evolutionary history of well-known proteins and pathways and potential novel functional diversity in animals.

## Acknowledgements

We thank members of the Snell-Rood Lab for critical feedback on this manuscript. We also thank the UMN University Imaging Centers for providing the training and equipment to obtain in situ images.

## Notes

### Competing Interest Statement

The authors have declared no competing interest.

## References

Altschul, S.F., Gish, W., Miller, W., Myers, E.W., Lipman, D.J., 1990. Basic local alignment search tool. J. Mol. Biol. 215, 403–410. 10.1016/S0022-2836(05)80360-2

Arendt, D., Tessmar-Raible, K., Snyman, H., Dorresteijn, A.W., Wittbrodf, J., 2004. Ciliary photoreceptors with a vertebrate-type opsin in an invertebrate brain. Science (80-.). 306, 869–871. 10.1126/science.1099955

Babu, Y.S., Bugg, C.E., Cook, W.J., 1988. Structure of calmodulin refined at 2.2 Å resolution. J. Mol. Biol. 204, 191–204. 10.1016/0022-2836(88)90608-0

Bej, A., Ames, J.B., 2022. Retinal Cyclic Nucleotide-Gated Channel Regulation by Calmodulin. Int. J. Mol. Sci. 23. 10.3390/ijms232214143

Burgoyne, R.D., Weiss, J.L., 2001. The neuronal calcium sensor family of Ca2+-binding proteins. Biochem. J. 353, 1–12. 10.1042/bj3530001

Chen, L., Li, G., Jiang, Z., Yau, K.-W., 2023. Unusual phototransduction via cross-motif signaling from G q to adenylyl cyclase in intrinsically photosensitive retinalganglion cells. Proc. Natl. Acad. Sci. 120, 2017. 10.1073/pnas.2216599120

Del Pilar Gomez, M.D.P., Nasi, E., 2000. Light transduction in invertebrate hyperpolarizing photoreceptors: Possible involvement of a G(o)-regulated guanylate cyclase. J. Neurosci. 20, 5254–5263. 10.1523/jneurosci.20-14-05254.2000

Dexter, P.M., Lobanova, E.S., Finkelstein, S., Spencer, W.J., Skiba, N.P., Arshavsky, V.Y., 2018. Transducin β-subunit can interact with multiple g protein γ-subunits to enable light detection by rod photoreceptors. eNeuro 5, 1–9. 10.1523/ENEURO.0144-18.2018

Dizhoor, A.M., Lowe, D.G., Olshevskaya, E. V., Laura, R.P., Hurley, J.B., 1994. The human photoreceptor membrane guanylyl cyclase, RetGC, is present in outer segments and is regulated by calcium and a soluble activator. Neuron 12, 1345–1352. 10.1016/0896-6273(94)90449-9

Döring, C.C., Kumar, S., Tumu, S.C., Kourtesis, I., Hausen, H., 2020. The visual pigment xenopsin is widespread in protostome eyes and impacts the view on eye evolution. Elife 9, 1–23. 10.7554/ELIFE.55193

Ekström, P., Garm, A., Pålsson, J., Vihtelic, T.S., Nilsson, D.E., 2008. Immunohistochemical evidence for multiple photosystems in box jellyfish. Cell Tissue Res. 333, 115–124. 10.1007/s00441-008-0614-8

Fain, G.L., Hardie, R., Laughlin, S.B., 2010. Phototransduction and the Evolution of Photoreceptors. Curr. Biol. 20, R114–R124. 10.1016/j.cub.2009.12.006

Faurobert, E., Chen, C.K., Hurley, J.B., Teng, D.H.F., 1996. Drosophila neurocalcin, a fatty acylated, Ca2+-binding protein that associates with membranes and inhibits in Vitro phosphorylation of bovine rhodopsin. J. Biol. Chem. 271, 10256–10262. 10.1074/jbc.271.17.10256

Firestein, S., 2001. How the olfactory system makes sense of scents. Nature 413, 211–218. 10.1038/35093026

Gold, D.A., Katsuki, T., Li, Y., Yan, X., Regulski, M., Ibberson, D., Holstein, T., Steele, R.E., Jacobs, D.K., Greenspan, R.J., 2019. The genome of the jellyfish Aurelia and the evolution of animal complexity. Nat. Ecol. Evol. 3, 96–104. 10.1038/s41559-018-0719-8

Gornik, S.G., Bergheim, B.G., Morel, B., Stamatakis, A., Foulkes, N.S., Guse, A., 2021. Photoreceptor Diversification Accompanies the Evolution of Anthozoa. Mol. Biol. Evol. 38, 1744–1760. 10.1093/molbev/msaa304

Gurevich, E. V., Gurevich, V. V., 2006. Arrestins: Ubiquitous regulators of cellular signaling pathways. Genome Biol. 7. 10.1186/gb-2006-7-9-236

Hardie, R.C., Raghu, P., 2001. Visual transduction in Drosophila. Nature. 10.1038/35093002

Hattar, S., Lucas, R.J., Mrosovsky, N., Thompson, S., Douglas, R.H., Hankins, M.W., Lem, J., Biel, M., Hofmann, F., Foster, R.G., Yau, K.W., 2003. Melanopsin and rod—cone photoreceptive systems account for all major accessory visual functions in mice. Nature 424, 76–81. 10.1038/nature01761

Ikura, M., 1996. Calcium binding and conformational response in EF-hand proteins. Trends Biochem. Sci. 21, 14–17. 10.1016/S0968-0004(06)80021-6

Katoh, K., Standley, D.M., 2013. MAFFT multiple sequence alignment software version 7: Improvements in performance and usability. Mol. Biol. Evol. 30, 772–780. 10.1093/molbev/mst010

Katz, B., Minke, B., 2009. Drosophila photoreceptors and signaling mechanisms. Front. Cell. Neurosci. 3, 2. 10.3389/neuro.03.002.2009

Kayal, E., Bentlage, B., Sabrina Pankey, M., Ohdera, A.H., Medina, M., Plachetzki, D.C., Collins, A.G., Ryan, J.F., 2018. Phylogenomics provides a robust topology of the major cnidarian lineages and insights on the origins of key organismal traits. BMC Evol. Biol. 18, 1–18. 10.1186/s12862-018-1142-0

Koch, K.W., Stryer, L., 1988. Highly cooperative feedback control of retinal rod guanylate cyclase by calcium ions. Nature 334, 64–66. 10.1038/334064a0

Kojima, D., Terakita, A., Ishikawa, T., Tsukahara, Y., Maeda, A., Shichida, Y., 1997. A novel G(o)-mediated phototransduction cascade in scallop visual cells. J. Biol. Chem. 272, 22979–22982. 10.1074/jbc.272.37.22979

Koyanagi, M., Takano, K., Tsukamoto, H., Ohtsu, K., Tokunaga, F., Terakita, A., 2008. Jellyfish vision starts with cAMP signaling mediated by opsin-Gs cascade. Proc. Natl. Acad. Sci. U. S. A. 105, 15576–15580. 10.1073/pnas.0806215105

Kozmik, Z., Ruzickova, J., Jonasova, K., Matsumoto, Y., Vopalensky, P., Kozmikova, I., Strnad, H., Kawamura, S., Piatigorsky, J., Paces, V., Vlcek, C., 2008. Assembly of the cnidarian camera-type eye from vertebrate-like components. Proc. Natl. Acad. Sci. 105, 8989–8993. 10.1073/pnas.0800388105

Krishnan, A., Mustafa, A., Almén, M.S., Fredriksson, R., Williams, M.J., Schiöth, H.B., 2015. Evolutionary hierarchy of vertebrate-like heterotrimeric G protein families. Mol. Phylogenet. Evol. 91, 27–40. 10.1016/j.ympev.2015.05.009

Lagman, D., Haines, H.J., Abalo, X.M., Larhammar, D., 2022. Ancient multiplicity in cyclic nucleotide-gated (CNG) cation channel repertoire was reduced in the ancestor of Olfactores before re-expansion by whole genome duplications in vertebrates. PLoS One 17, 1–27. 10.1371/journal.pone.0279548

Lagman, D., Sundström, G., Daza, D.O., Abalo, X.M., Larhammar, D., 2012. Expansion of transducin subunit gene families in early vertebrate tetraploidizations. Genomics 100, 203–211. 10.1016/j.ygeno.2012.07.005

Lamb, T.D., 2013. Evolution of phototransduction, vertebrate photoreceptors and retina. Prog. Retin. Eye Res. 36, 52–119. 10.1016/j.preteyeres.2013.06.001

Lamb, T.D., Hunt, D.M., 2017. Evolution of the vertebrate phototransduction cascade activation steps. Dev. Biol. 431, 77–92. 10.1016/j.ydbio.2017.03.018

Lamb, T.D., Patel, H.R., Chuah, A., Hunt, D.M., 2018. Evolution of the shut-off steps of vertebrate phototransduction. Open Biol. 8, 170232. 10.1098/rsob.170232

Layden, M.J., Rentzsch, F., Röttinger, E., 2016. The rise of the starlet sea anemone Nematostella vectensis as a model system to investigate development and regeneration. Wiley Interdiscip. Rev. Dev. Biol. 5, 408–428. 10.1002/wdev.222

Lee, S.J., Xu, H., Montell, C., 2004. Rhodopsin kinase activity modulates the amplitude of the visual response in Drosophila. Proc. Natl. Acad. Sci. U. S. A. 101, 11874–11879. 10.1073/pnas.0402205101

Letunic, I., Bork, P., 2021. Interactive tree of life (iTOL) v5: An online tool for phylogenetic tree display and annotation. Nucleic Acids Res. 49, W293–W296. 10.1093/nar/gkab301

Levay, K., Cabrera, J.L., Satpaev, D.K., Slepak, V.Z., 1999. Gβ5 prevents the RGS7-Gαo interaction through binding to a distinct Gγ-like domain found in RGS7 and other RGS proteins. Proc. Natl. Acad. Sci. U. S. A. 96, 2503–2507. 10.1073/pnas.96.5.2503

Liegertová, M., Pergner, J., Kozmiková, I., Fabian, P., Pombinho, A.R., Strnad, H., Pačes, J., Vlček, Č., Bartůněk, P., Kozmik, Z., 2015. Cubozoan genome illuminates functional diversification of opsins and photoreceptor evolution. Sci. Rep. 5, 1–19. 10.1038/srep11885

Lokits, A.D., Indrischek, H., Meiler, J., Hamm, H.E., Stadler, P.F., 2018. Tracing the evolution of the heterotrimeric G protein α subunit in Metazoa. BMC Evol. Biol. 18, 1–27. 10.1186/s12862-018-1147-8

Lyubarsky, A.L., Chen, C.K., Simon, M.I., Pugh, E.N., 2000. Mice lacking G-protein receptor kinase 1 have profoundly slowed recovery of cone-driven retinal responses. J. Neurosci. 20, 2209–2217. 10.1523/jneurosci.20-06-02209.2000

Macias-Munõz, A., Murad, R., Mortazavi, A., 2019. Molecular evolution and expression of opsin genes in Hydra vulgaris. BMC Genomics 20, 1–19. 10.1186/s12864-019-6349-y

Makino, E.R., Handy, J.W., Li, T., Arshavsky, V.Y., 1999. The GTPase activating factor for transducin in rod photoreceptors is the complex between RGS9 and type 5 G protein β subunit. Proc. Natl. Acad. Sci. U. S. A. 96, 1947–1952. 10.1073/pnas.96.5.1947

Mason, B., Schmale, M., Gibbs, P., Miller, M.W., Wang, Q., Levay, K., Shestopalov, V., Slepak, V.Z., 2012. Evidence for Multiple Phototransduction Pathways in a Reef-Building Coral. PLoS One 7, 1–9. 10.1371/journal.pone.0050371

McCulloch, K., Liu, A., Babonis, L., Daly, C., Martindale, M.Q., Koenig, K., 2023. *Nematostella vectensis* exemplifies the exceptional expansion and diversity of opsins in the eyeless Hexacorallia. bioRxiv 2023.05.17. 10.1101/2023.05.17.541201

Minh, B.Q., Schmidt, H.A., Chernomor, O., Schrempf, D., Woodhams, M.D., Von Haeseler, A., Lanfear, R., Teeling, E., 2020. IQ-TREE 2: New Models and Efficient Methods for Phylogenetic Inference in the Genomic Era. Mol. Biol. Evol. 37, 1530–1534. 10.1093/molbev/msaa015

Montell, C., 2005. TRP channels in Drosophila photoreceptor cells. J. Physiol. 567, 45–51. 10.1113/jphysiol.2005.092551

Moroz, L.L., Mukherjee, K., Romanova, D.Y., 2023. Nitric oxide signaling in ctenophores. Front. Neurosci. 17, 1–15. 10.3389/fnins.2023.1125433

Ohdera, A., Ames, C.L., Dikow, R.B., Kayal, E., Chiodin, M., Busby, B., La, S., Pirro, S., Collins, A.G., Medina, M., Ryan, J.F., 2019. Box, stalked, and upside-down? Draft genomes from diverse jellyfish (cnidaria, acraspeda) lineages: Alatina alata (cubozoa), calvadosia cruxmelitensis (staurozoa), and cassiopea xamachana (scyphozoa). Gigascience 8, 1–15. 10.1093/gigascience/giz069

Okada, T., Sugihara, M., Bondar, A.N., Elstner, M., Entel, P., Buss, V., 2004. The retinal conformation and its environment in rhodopsin in light of a new 2.2 Å crystal structure. J. Mol. Biol. 342, 571–583. 10.1016/j.jmb.2004.07.044

Oldham, W.M., Hamm, H.E., 2006. Structural basis of function in heterotrimeric G proteins. Q. Rev. Biophys. 39, 117–166. 10.1017/S0033583506004306

Palczewski, K., Kumasaka, T., Hori, T., Behnke, C.A., Motoshima, H., Fox, B.A., Trong, I. Le, Teller, D.C., Okada, T., Stenkamp, R.E., Yamamoto, M., Miyano, M., 2000. Crystal Structure of Rhodopsin: A G Protein-Coupled Receptor. Science 289, 739–745. 10.1126/science.289.5480.739

Paysan-Lafosse, T., Blum, M., Chuguransky, S., Grego, T., Pinto, B.L., Salazar, G.A., Bileschi, M.L., Bork, P., Bridge, A., Colwell, L., Gough, J., Haft, D.H., Letunić, I., Marchler-Bauer, A., Mi, H., Natale, D.A., Orengo, C.A., Pandurangan, A.P., Rivoire, C., Sigrist, C.J.A., Sillitoe, I., Thanki, N., Thomas, P.D., Tosatto, S.C.E., Wu, C.H., Bateman, A., 2023. InterPro in 2022. Nucleic Acids Res. 51, D418–D427. 10.1093/nar/gkac993

Peng, G., Shi, X., Kadowaki, T., 2015. Evolution of TRP channels inferred by their classification in diverse animal species. Mol. Phylogenet. Evol. 84, 145–157. 10.1016/j.ympev.2014.06.016

Peng, Y.W., Robishaw, J.D., Levine, M.A., Yau, K.W., 1992. Retinal rods and cones have distinct G protein β and γ subunits. Proc. Natl. Acad. Sci. U. S. A. 89, 10882–10886. 10.1073/pnas.89.22.10882

Plachetzki, D.C., Fong, C.R., Oakley, T.H., 2012. Cnidocyte discharge is regulated by light and opsin-mediated phototransduction. BMC Biol. 10. 10.1186/1741-7007-10-17

Plachetzki, D.C., Fong, C.R., Oakley, T.H., 2010. The evolution of phototransduction from an ancestral cyclic nucleotide gated pathway. Proc. R. Soc. B Biol. Sci. 277, 1963–1969. 10.1098/rspb.2009.1797

Potter, L.R., 2011. Guanylyl cyclase structure, function and regulation. Cell. Signal. 23, 1921–1926. 10.1016/j.cellsig.2011.09.001

Premont, R.T., Koch, W.J., Inglese, J., Lefkowitz, R.J., 1994. Identification, purification, and characterization of GRK5, a member of the family of G protein-coupled receptor kinases. J. Biol. Chem. 269, 6832–6841. 10.1016/s0021-9258(17)37451-3

Price, M.N., Dehal, P.S., Arkin, A.P., 2010. FastTree 2 - Approximately maximum-likelihood trees for large alignments. PLoS One 5. 10.1371/journal.pone.0009490

Ramirez, M.D., Pairett, A.N., Pankey, M.S., Serb, J.M., Speiser, D.I., Swafford, A.J., Oakley, T.H., 2016. The last common ancestor of most bilaterian animals possessed at least nine opsins. Genome Biol. Evol. 8, 3640–3652. 10.1093/gbe/evw248

Rawlinson, K.A., Lapraz, F., Ballister, E.R., Terasaki, M., Rodgers, J., McDowell, R.J., Girstmair, J., Criswell, K.E., Boldogkoi, M., Simpson, F., Goulding, D., Cormie, C., Hall, B., Lucas, R.J., Telford, M.J., 2019. Extraocular, rod-like photoreceptors in a flatworm express xenopsin photopigment. Elife 8, 1–28. 10.7554/eLife.45465

Schnitzler, C.E., Pang, K., Powers, M.L., Reitzel, A.M., Ryan, J.F., Simmons, D., Tada, T., Park, M., Gupta, J., Brooks, S.Y., Blakesley, R.W., Yokoyama, S., Haddock, S.H.D., Martindale, M.Q., Baxevanis, A.D., 2012. Genomic organization, evolution, and expression of photoprotein and opsin genes in Mnemiopsis leidyi: A new view of ctenophore photocytes. BMC Biol. 10. 10.1186/1741-7007-10-107

Sebé-Pedrós, A., Saudemont, B., Chomsky, E., Plessier, F., Mailhé, M.P., Renno, J., Loe-Mie, Y., Lifshitz, A., Mukamel, Z., Schmutz, S., Novault, S., Steinmetz, P.R.H., Spitz, F., Tanay, A., Marlow, H., 2018. Cnidarian Cell Type Diversity and Regulation Revealed by Whole-Organism Single-Cell RNA-Seq. Cell 173, 1520–1534.e20. 10.1016/j.cell.2018.05.019

Shichida, Y., Matsuyama, T., 2009. Evolution of opsins and phototransduction. Philos. Trans. R. Soc. B Biol. Sci. 364, 2881–2895. 10.1098/rstb.2009.0051

Simon, M.I., Strathmann, M.P., Gautam, N., 1991. Diversity of G proteins in signal transduction. Science (80-.). 252, 802–808. 10.1126/science.1902986

Squires, K.E., Montañez-Miranda, C., Pandya, R.R., Torres, M.P., Hepler, J.R., 2018. Genetic analysis of rare human variants of regulators of G protein signaling proteins and their role in human physiology and disease. Pharmacol. Rev. 70, 446–474. 10.1124/pr.117.015354

Suga, H., Schmid, V., Gehring, W.J., 2008. Evolution and Functional Diversity of Jellyfish Opsins. Curr. Biol. 18, 51–55. 10.1016/j.cub.2007.11.059

Sun, Z.L., Zheng, Y.H., Liu, W., 2018. Identification and characterization of a novel calmodulin binding site in Drosophila TRP C-terminus. Biochem. Biophys. Res. Commun. 501, 434–439. 10.1016/j.bbrc.2018.05.007

Swardfager, W., Mitchell, J., 2007. Purification of visual arrestin from squid photoreceptors and characterization of arrestin interaction with rhodopsin and rhodopsin kinase. J. Neurochem. 101, 223–231. 10.1111/j.1471-4159.2006.04364.x

Tanaka, T., Ames, J.B., Harvey, T.S., Stryer, L., Ikura, M., 1995. Sequestration of the membrane-targeting myristoyl group of recoverin in the calcium-free state. Nature 376, 444–447. 10.1038/376444a0

Tarrant, A.M., Helm, R.R., Levy, O., Rivera, H.E., 2019. Environmental entrainment demonstrates natural circadian rhythmicity in the cnidarian Nematostella vectensis. J. Exp. Biol. 222. 10.1242/jeb.205393

Terakita, A., 2005. The opsins. Genome Biol. 6, 1–9. 10.1186/gb-2005-6-3-213

Trudeau, M.C., Zagotta, W.N., 2004. Dynamics of Ca2+-calmodulin-dependent inhibition of rod cyclic nucleotide-gated channels measured by patch-clamp fluorometry. J. Gen. Physiol. 124, 211–223. 10.1085/jgp.200409101

Velarde, R.A., Sauer, C.D., Walden, K.K.O., Fahrbach, S.E., Robertson, H.M., 2005. Pteropsin: A vertebrate-like non-visual opsin expressed in the honey bee brain. Insect Biochem. Mol. Biol. 35, 1367–1377. 10.1016/j.ibmb.2005.09.001

Vöcking, O., Kourtesis, I., Tumu, S.C., Hausen, H., 2017. Co-expression of xenopsin and rhabdomeric opsin in photoreceptors bearing microvilli and cilia. Elife 6, 1–26. 10.7554/eLife.23435

Vöcking, O., Macias-Muñoz, A., Jaeger, S.J., Oakley, T.H., 2022. Deep Diversity: Extensive Variation in the Components of Complex Visual Systems across Animals. Cells 11, 3966. 10.3390/cells11243966

Wang, T., Montell, C., 2007. Phototransduction and retinal degeneration in Drosophila. Pflugers Arch. Eur. J. Physiol. 454, 821–847. 10.1007/s00424-007-0251-1

Warner, J.F., Guerlais, V., Amiel, A.R., Johnston, H., Nedoncelle, K., Röttinger, E., 2018. NvERTx: a gene expression database to compare embryogenesis and regeneration in the sea anemone *Nematostella vectensis*. Development 145, dev162867. 10.1242/dev.162867

Wilden, U., Hall, S.W., Kuhn, H., 1986. Phosphodiesterase activation by photoexcited rhodopsin is quenched when rhodopsin is phosphorylated and binds the intrinsic 48-kDa protein of rod outer segments. Proc. Natl. Acad. Sci. U. S. A. 83, 1174–1178. 10.1073/pnas.83.5.1174

Yamashita, T., Ohuchi, H., Tomonari, S., Ikeda, K., Sakai, K., Shichida, Y., 2010. Opn5 is a UV-sensitive bistable pigment that couples with Gi subtype of G protein. Proc. Natl. Acad. Sci. U. S. A. 107, 22084–22089. 10.1073/pnas.1012498107

Yau, K.W., 1994. Phototransduction mechanism in retinal rods and cones. The Friedenwald lecture. Investig. Ophthalmol. Vis. Sci. 35, 9–32.

Yau, K.W., Hardie, R.C., 2009. Phototransduction Motifs and Variations. Cell 139, 246–264. 10.1016/j.cell.2009.09.029

Zimmermann, B., Robb, S.M.C., Genikhovich, G., Fropf, W.J., Weilguny, L., He, S., Chen, S., Lovegrove-Walsh, J., Hill, E.M., Ragkousi, K., Praher, D., Fredman, D., Moran, Y., Gibson, M.C., Technau, U., 2020. Sea anemone genomes reveal ancestral metazoan chromosomal macrosynteny. bioRxiv 2020.10.30.359448. 10.1101/2020.10.30.359448

