## Supplemental Figures for "Multiple parallel expansions of bilaterian-like phototransduction gene families in the eyeless Anthozoa"

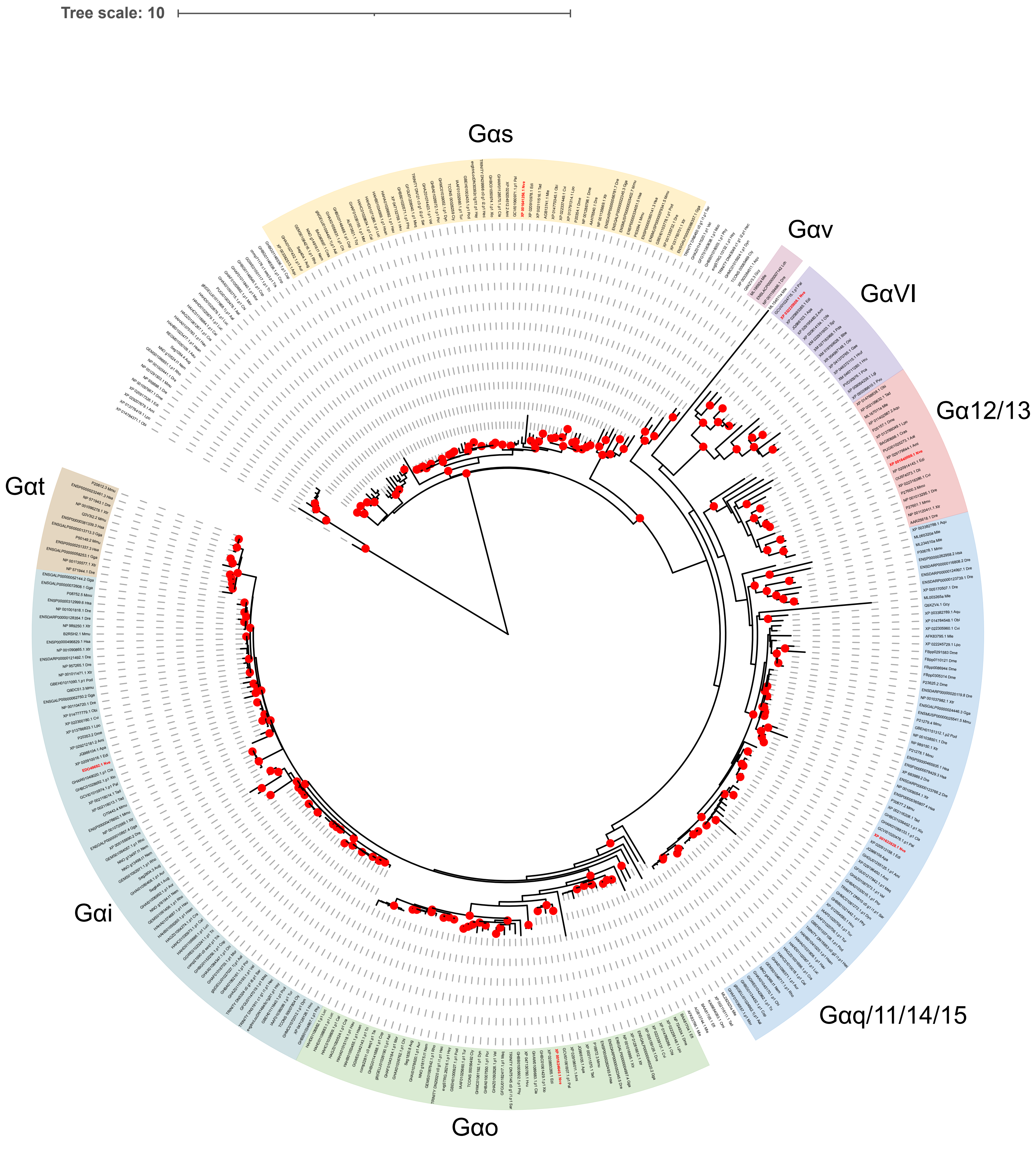


**Figure S1.** Maximum-likelihood tree of G alpha subunit families. Red circles at nodes indicate highly supported branch leading to that node, with 80% aSH-LRT/95% UFbs support. *Nematostella vectensis* sequences are bolded in red text.

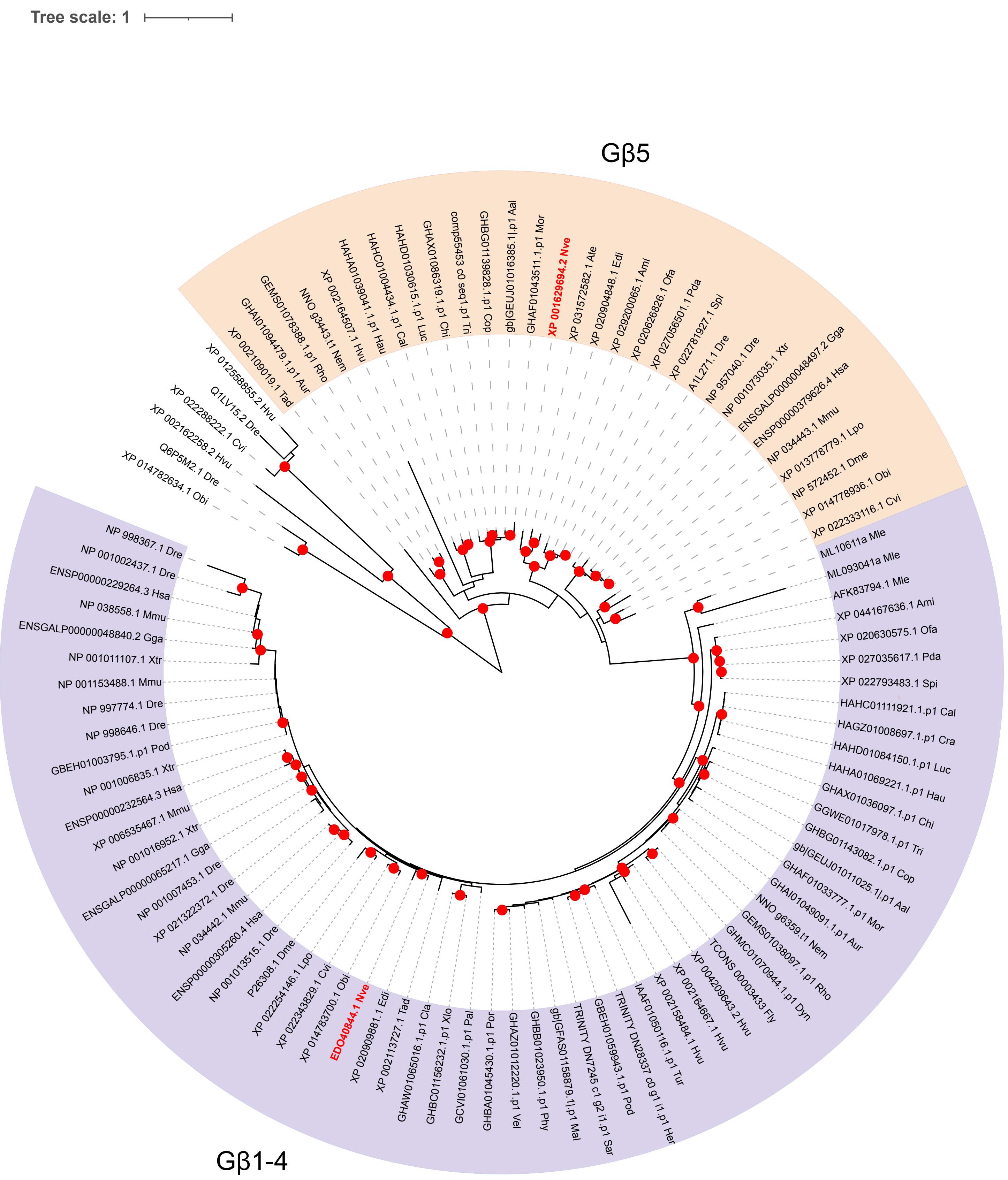


**Figure S2.** Maximum-likelihood tree of G protein beta subunit families. Red circles at nodes indicate highly supported branch leading to that node, with 80% aSH-LRT/95% UFbs support. *Nematostella vectensis* sequences are bolded in red text.

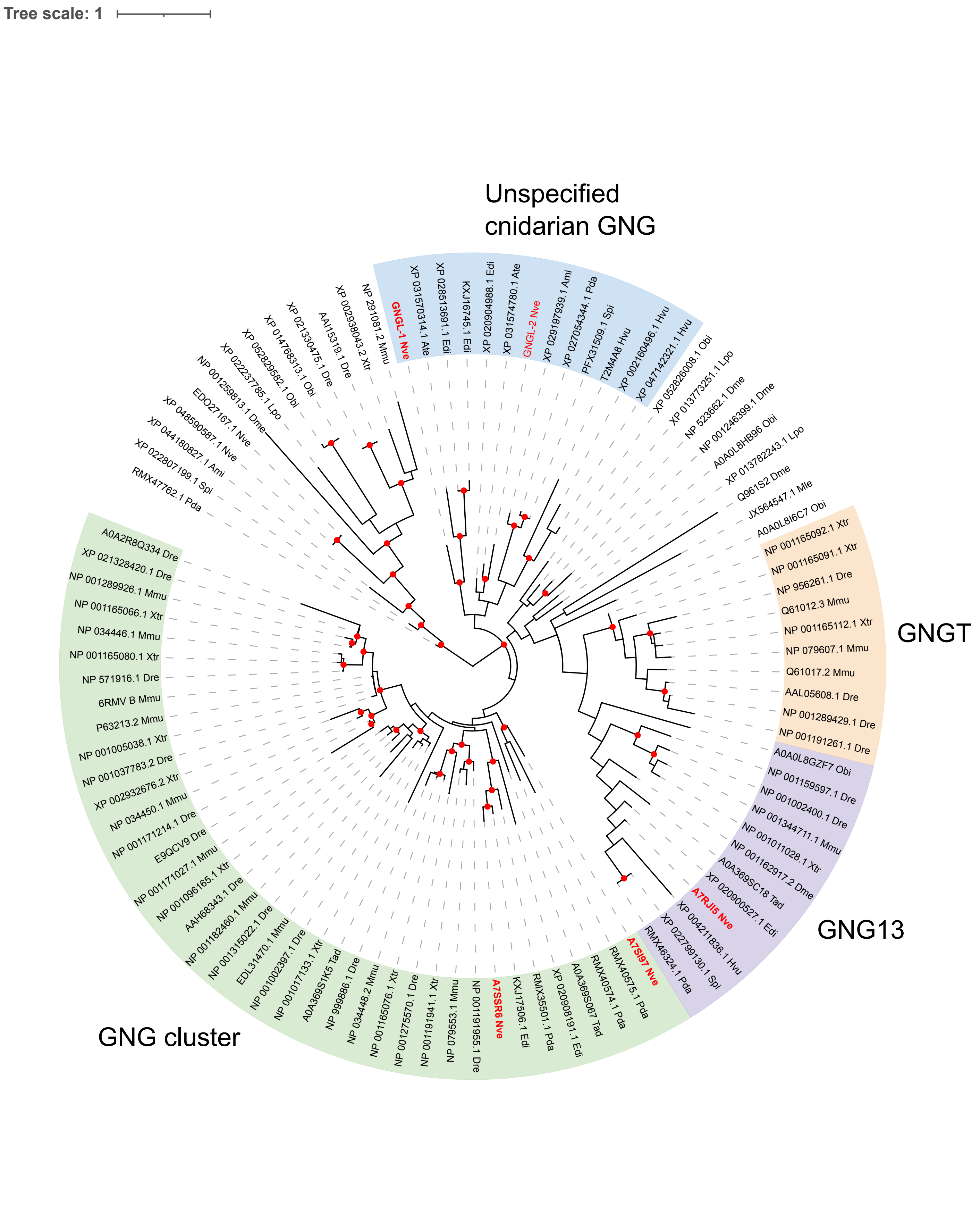


**Figure S3.** Maximum-likelihood tree of G gamma subunit families. Red circles at nodes indicate highly supported branch leading to that node, with 80% aSH-LRT/95% UFbs support. *Nematostella vectensis* sequences are bolded in red text.

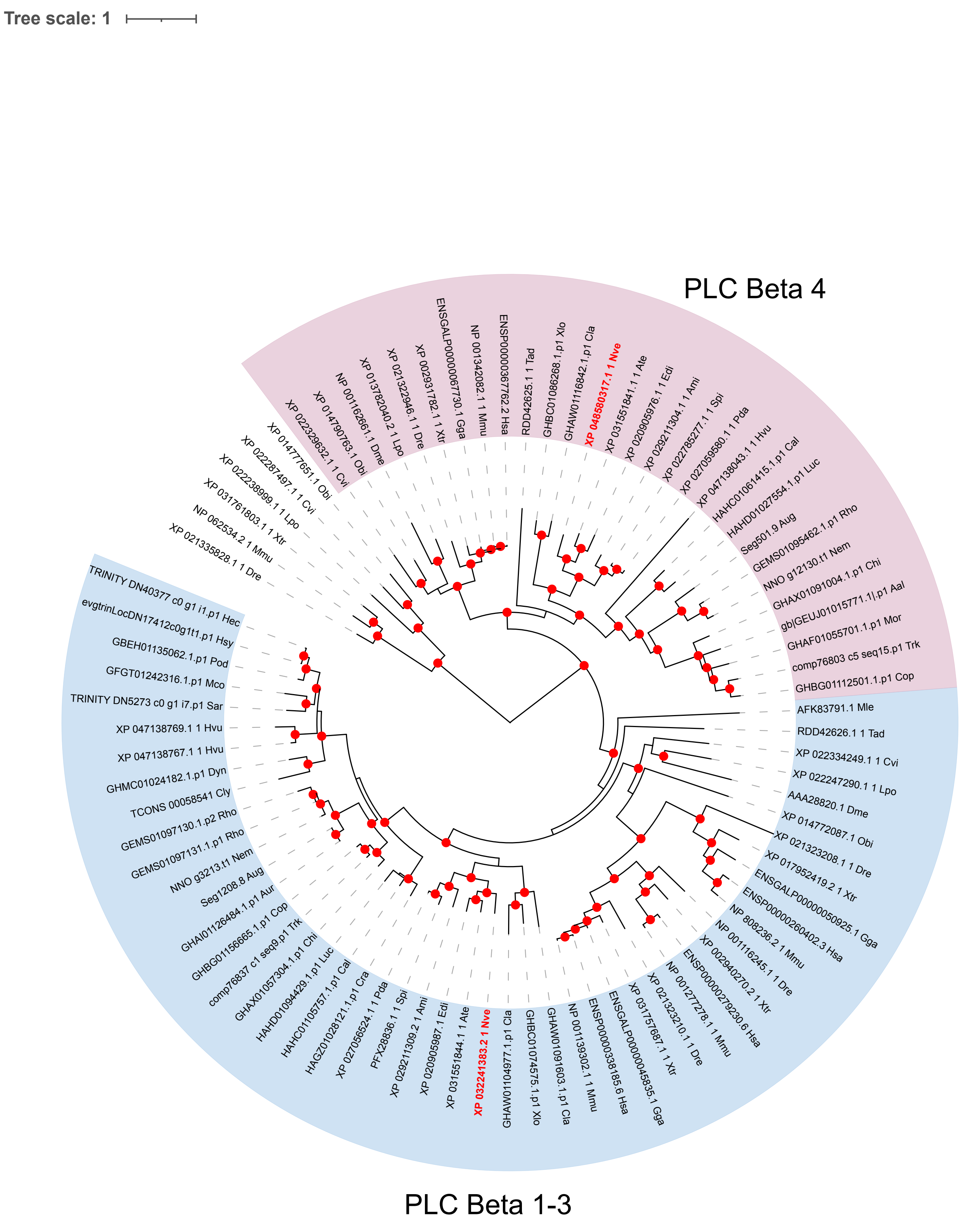


**Figure S4.** Maximum-likelihood tree of PLC beta. Red circles at nodes indicate highly supported branch leading to that node, with 80% aSH-LRT/95% UFbs support. *Nematostella vectensis* sequences are bolded in red text.

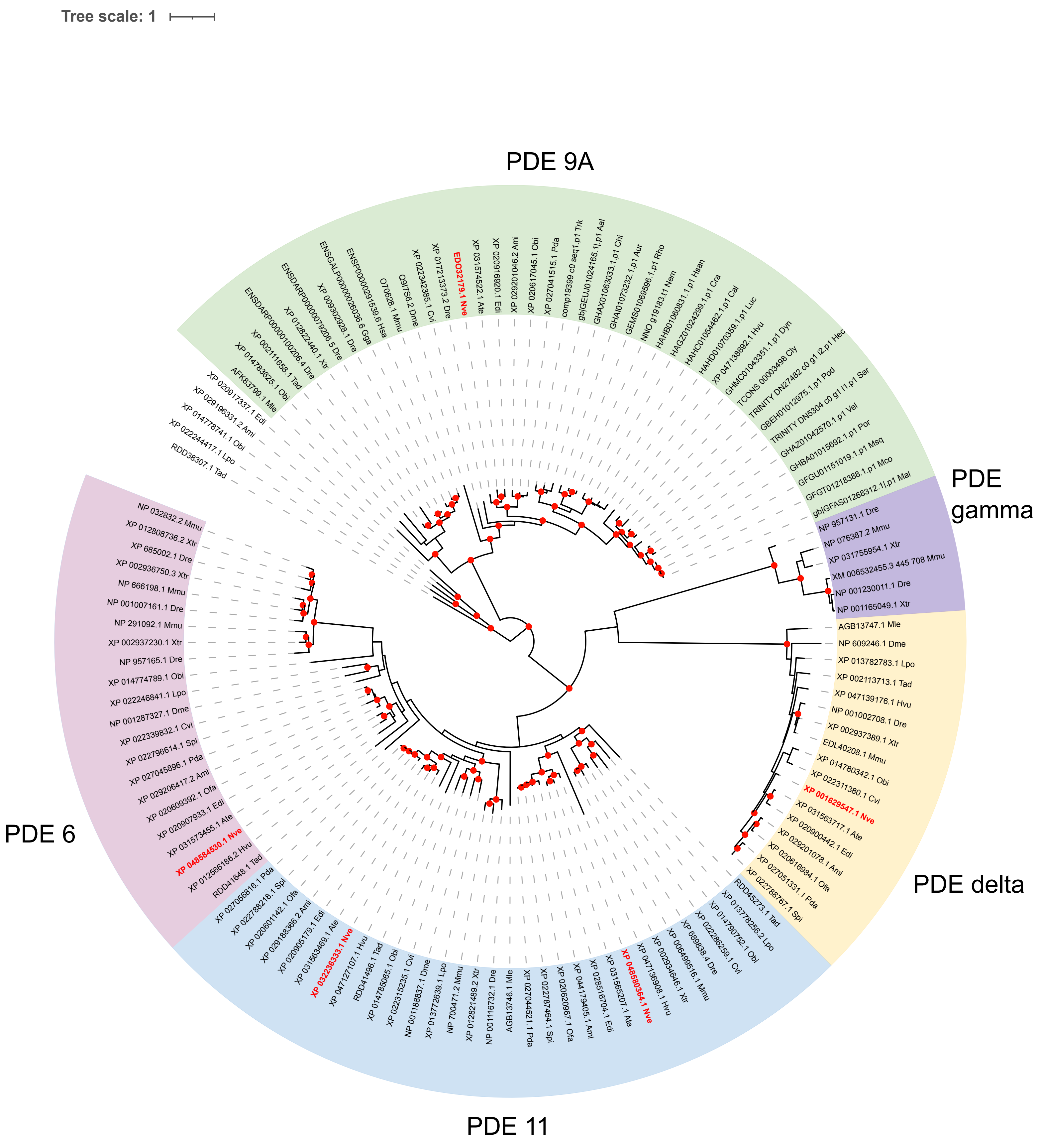


**Figure S5.** Maximum-likelihood tree of cGMP PDE subfamilies. Red circles at nodes indicate highly supported branch leading to that node, with 80% aSH-LRT/95% UFbs support. *Nematostella vectensis* sequences are bolded in red text.

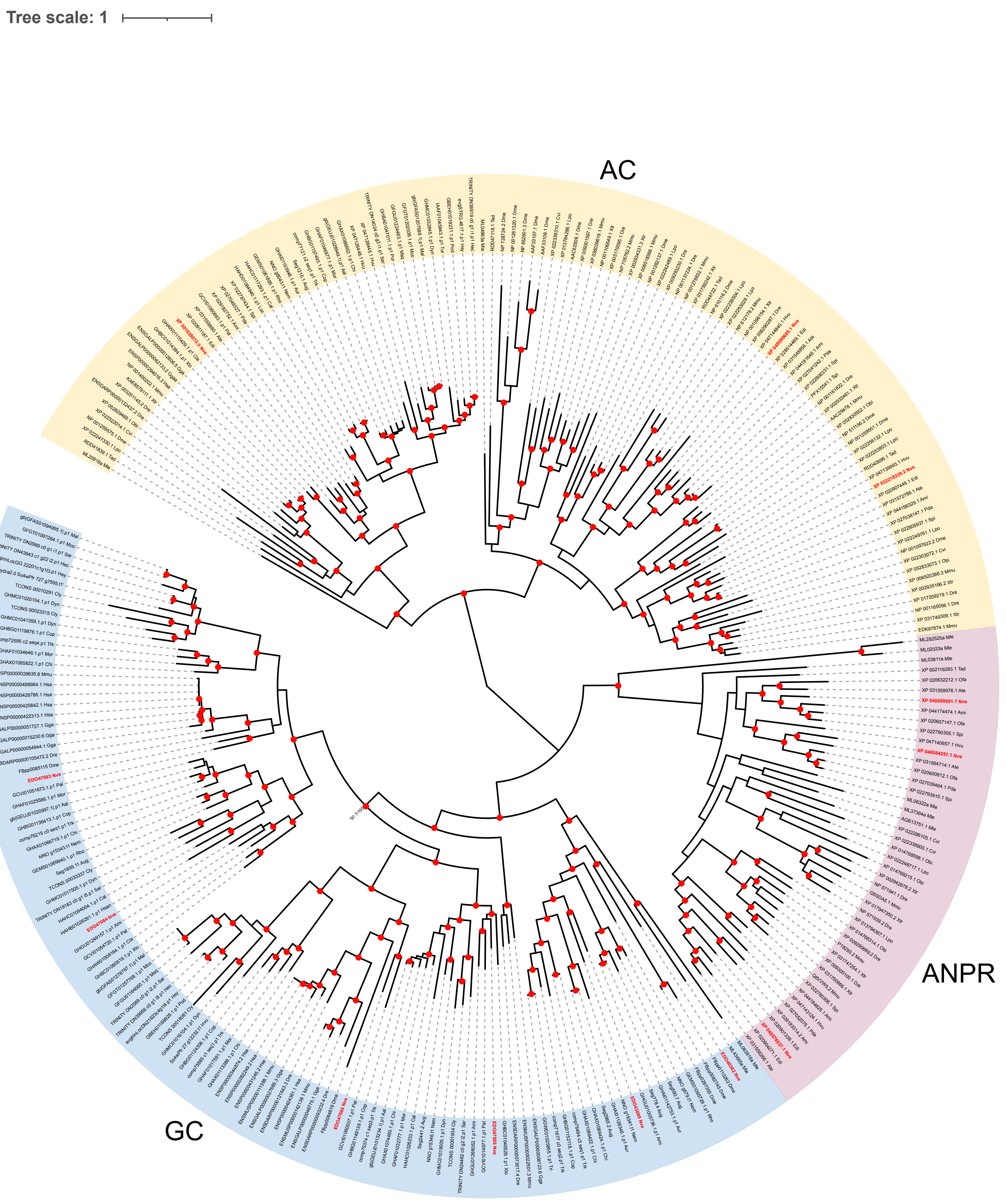


**Figure S6.** Maximum-likelihood tree of ANPR/GC and AC families. Red circles at nodes indicate highly supported branch leading to that node, with 80% aSH-LRT/95% UFbs support. *Nematostella vectensis* sequences are bolded in red text.

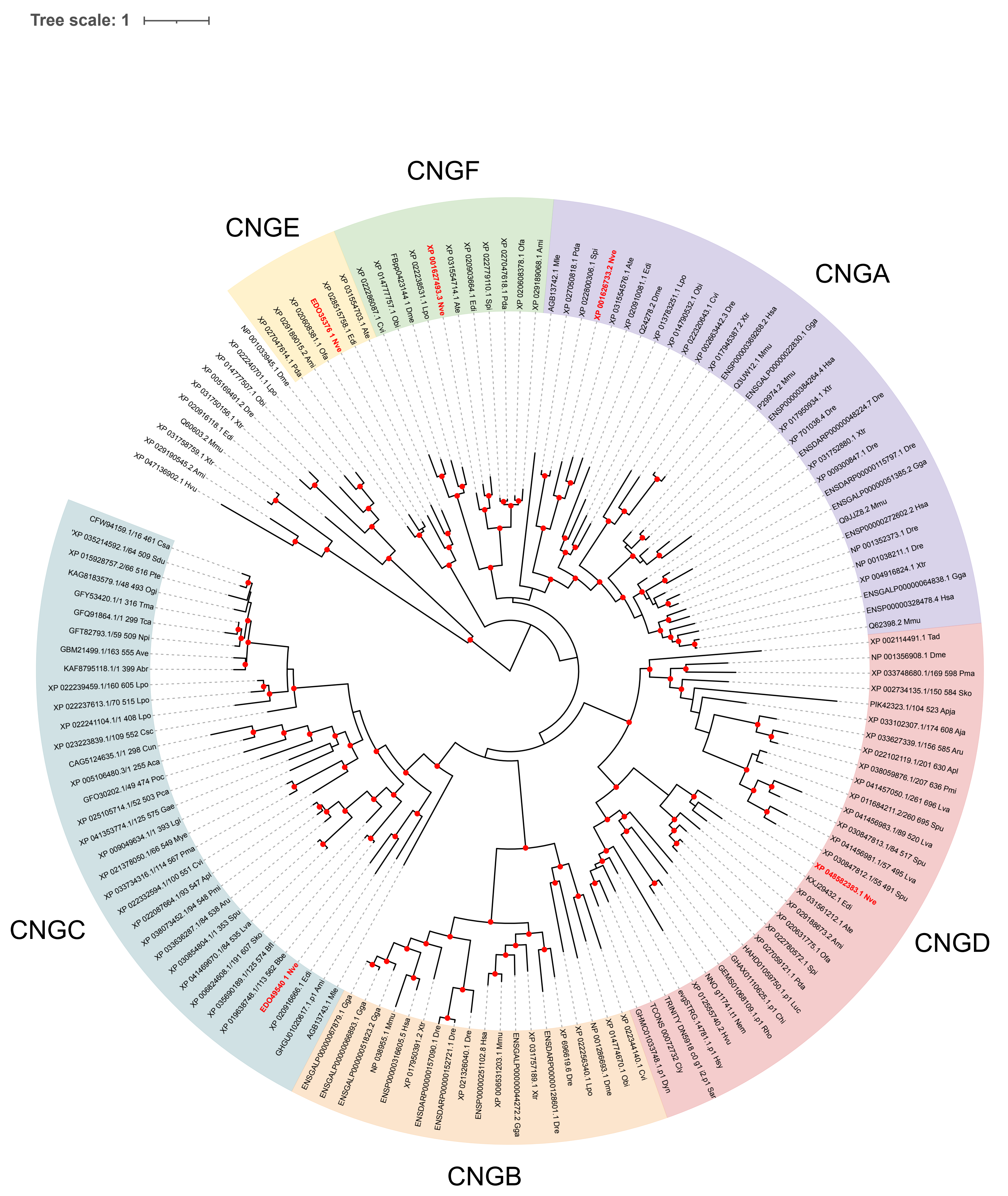


**Figure S7.** Maximum-likelihood tree of CNG channels. Red circles at nodes indicate highly supported branch leading to that node, with 80% aSH-LRT/95% UFbs support. *Nematostella vectensis* sequences are bolded in red text.

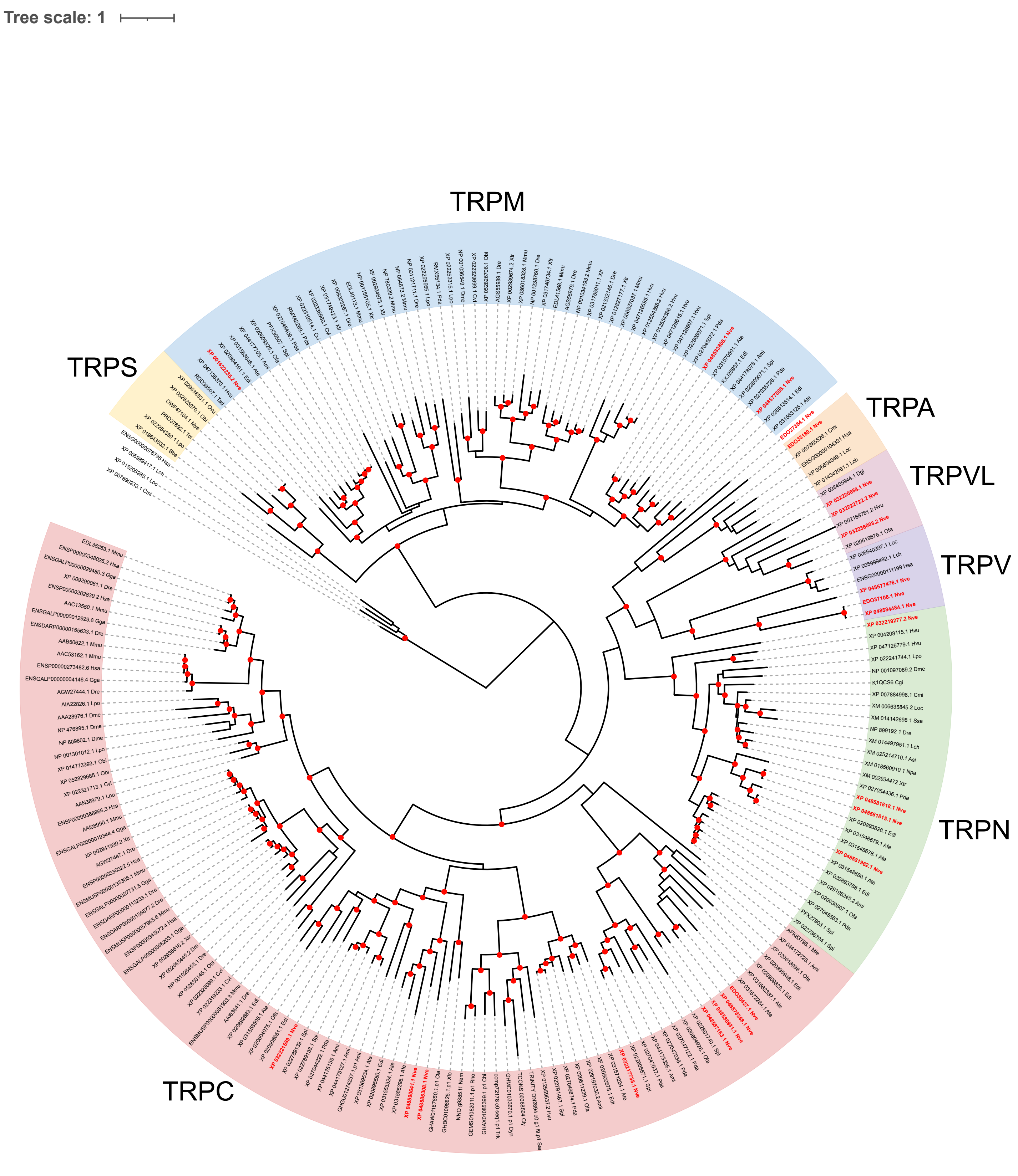


**Figure S8.** Maximum-likelihood tree of TRP channel families. Red circles at nodes indicate highly supported branch leading to that node, with 80% aSH-LRT/95% UFbs support. *Nematostella vectensis* sequences are bolded in red text.

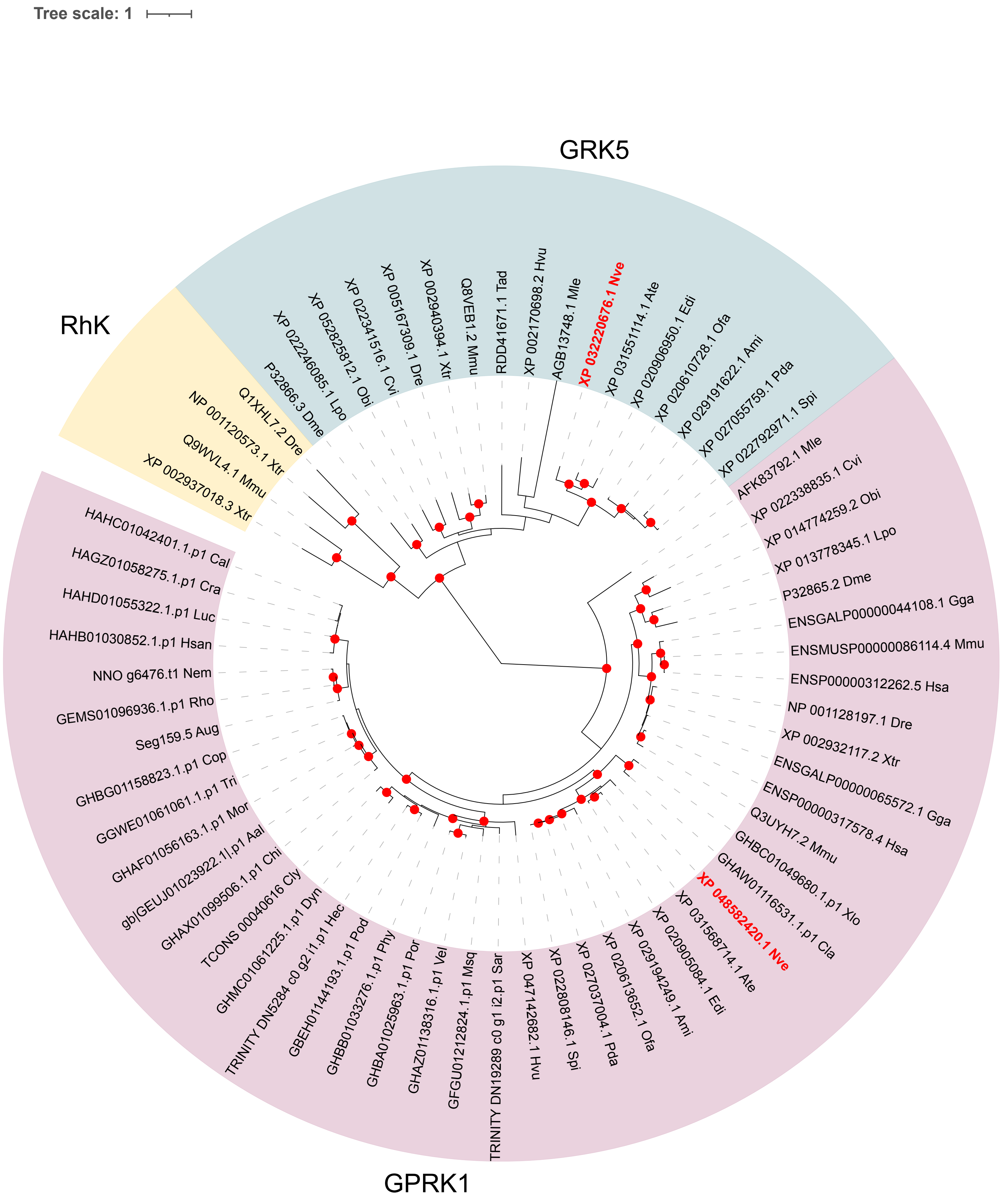


**Figure S9.** Maximum-likelihood tree of some G protein receptor kinases. Red circles at nodes indicate highly supported branch leading to that node, with 80% aSH-LRT/95% UFbs support. *Nematostella vectensis* sequences are bolded in red text.

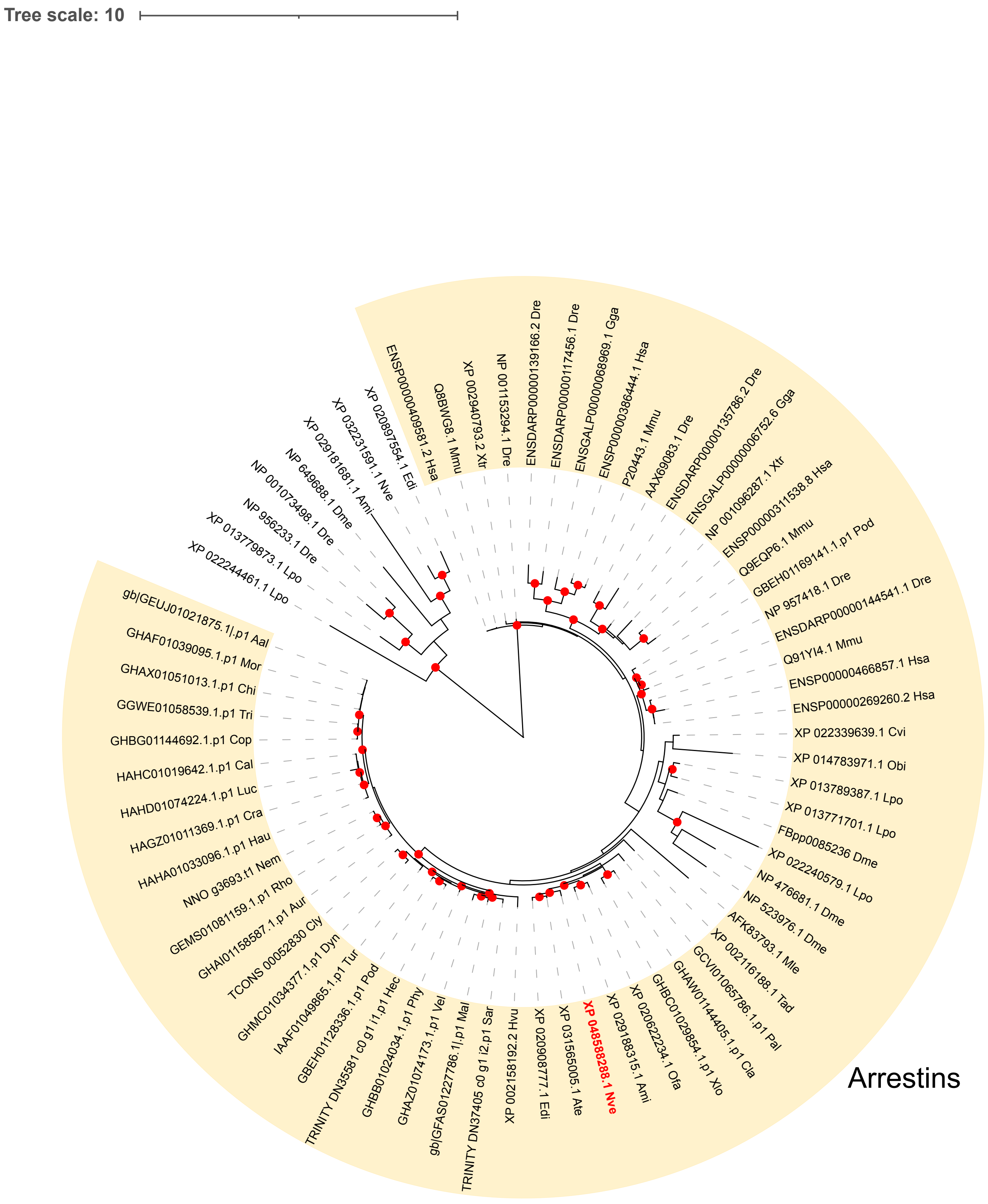


**Figure S10.** Maximum-likelihood tree of visual arrestin family. Red circles at nodes indicate highly supported branch leading to that node, with 80% aSH-LRT/95% UFbs support. *Nematostella vectensis* sequences are bolded in red text.

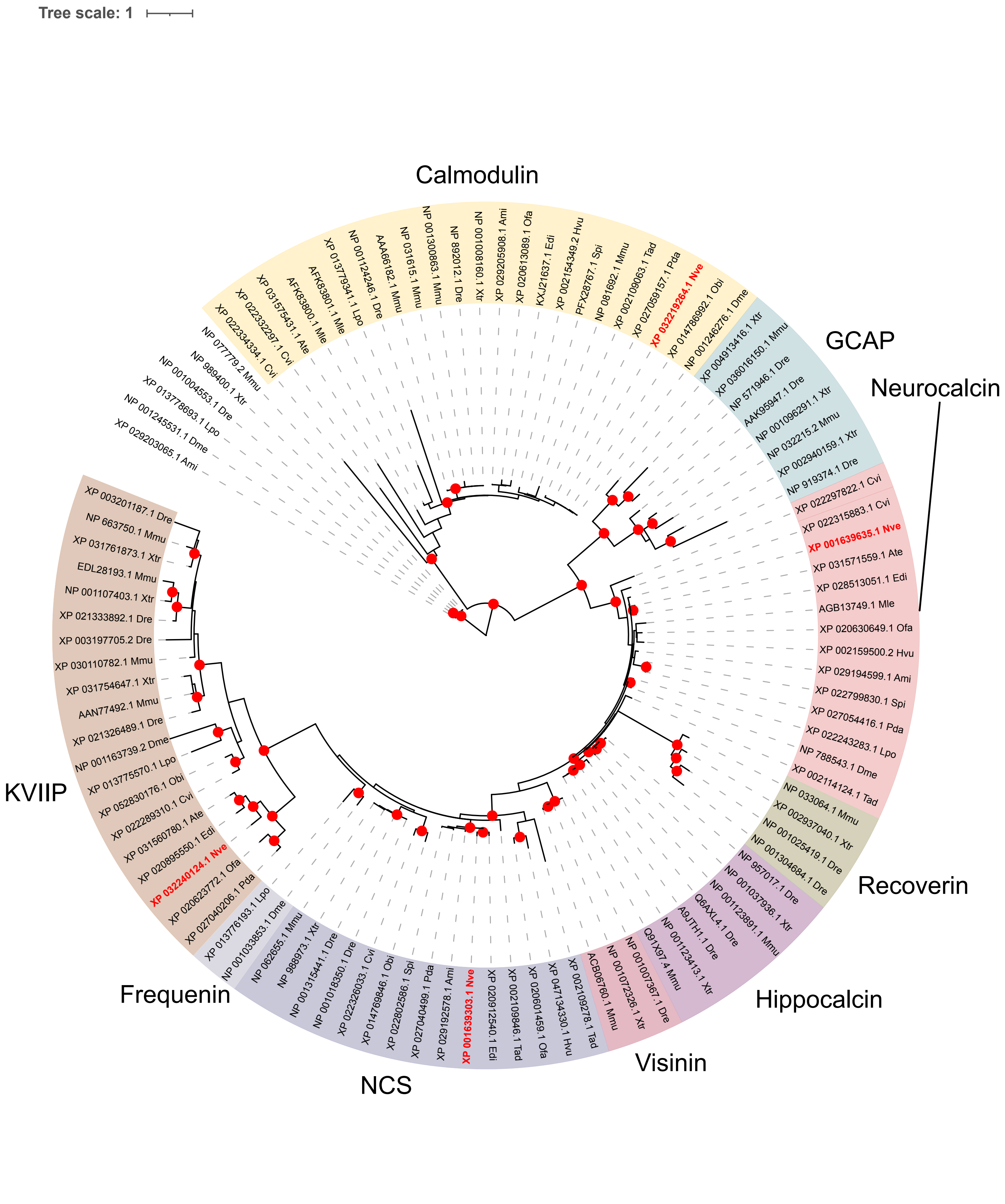


**Figure S11.** Maximum-likelihood tree of Calcium binding protein families. Red circles at nodes indicate highly supported branch leading to that node, with 80% aSH-LRT/95% UFbs support. *Nematostella vectensis* sequences are bolded in red text.

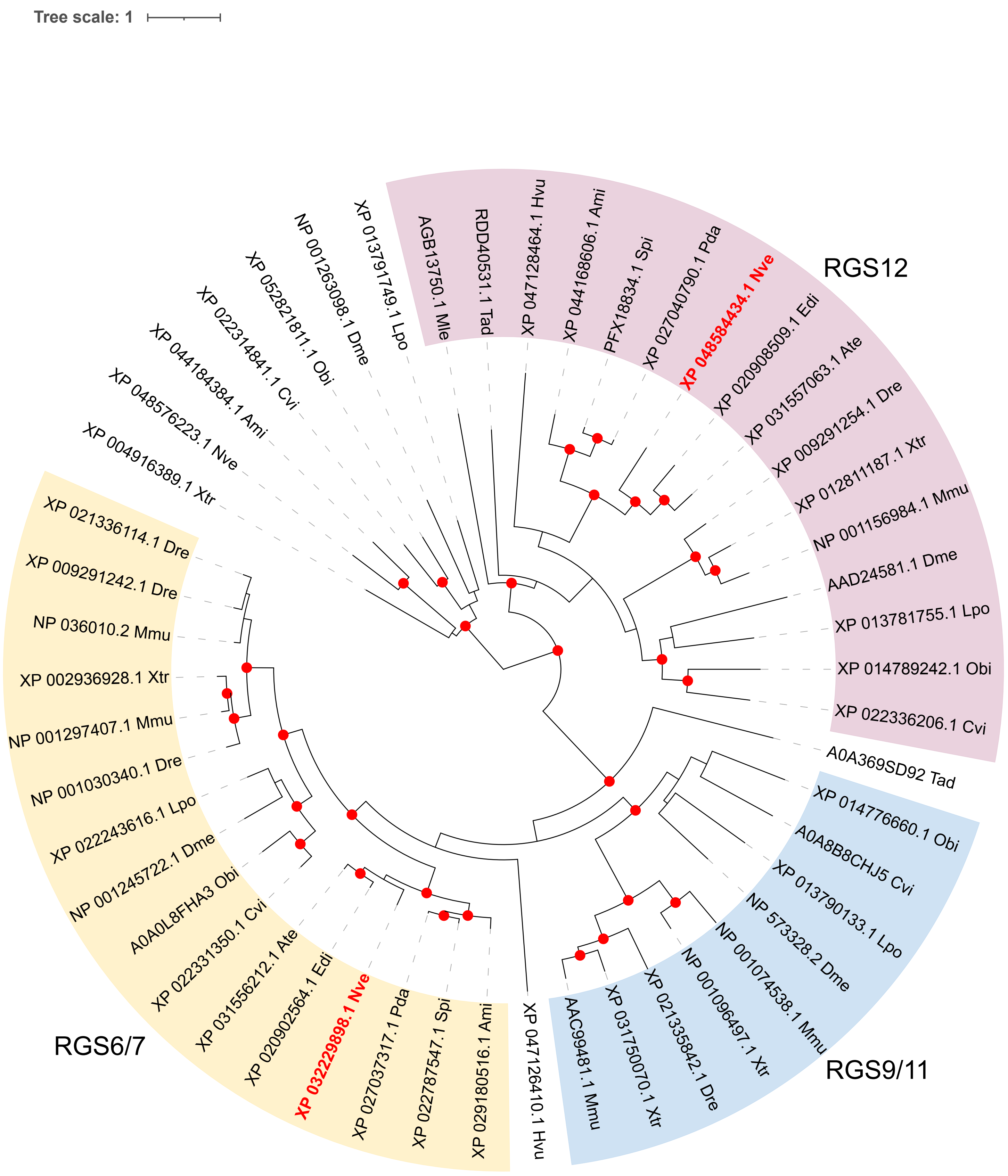


**Figure S12.** Maximum-likelihood tree of RGS proteins. Red circles at nodes indicate highly supported branch leading to that node, with 80% aSH-LRT/95% UFbs support. *Nematostella vectensis* sequences are bolded in red text.
